# Identification of three apicoplast transporters with crucial roles in the malaria parasite life cycle

**DOI:** 10.64898/2026.11.24.768832

**Authors:** Laura J. Akkerman, Roos Weeda, Hester Vogelaar, Ezra T. Bekkering, Silvia Tassan-Lugrezin, Sebastian Buss, Eric Beitz, Laura E. de Vries, Taco W.A. Kooij

## Abstract

Malaria parasites depend on an intriguing endosymbiotic organelle, the apicoplast, for survival throughout the life cycle. The apicoplast is enclosed by four membranes and transport of metabolites across these membranes remains poorly understood. Thus far, only twelve (putative) membrane transport proteins have been confirmed to localize to the organelle. We report a crucial function in the apicoplast of three previously uncharacterized members of the major facilitator superfamily of transporters in *Plasmodium falciparum*. *MCP1* and *UMF* are essential for asexual blood-stage parasites and could only be deleted in the presence of an alternative pathway for producing isopentenyl pyrophosphate. Deletion of either gene resulted in disruption of apicoplast morphology and loss of the organellar genome, suggesting that both are required for apicoplast housekeeping. In contrast, *MCP2* was dispensable in the blood stages yet essential for parasite development in the mosquito. Oocysts in the mosquito midgut lacking *MCP2* were significantly smaller than wild-type oocysts and produced no sporozoites, similar to when apicoplast fatty acid metabolism is disrupted. Differences in apicoplast morphology became apparent five days after infection, with *MCP2*-deficient oocyst apicoplasts displaying a less extended, sometimes fragmented, network with thinner branches. Taken together, we identified three novel apicoplast transporters, with *MCP1* and *UMF* being essential for asexual blood-stage development by supporting apicoplast housekeeping, and *MCP2* being crucial for parasite growth in mosquitoes, possibly by facilitating fatty acid metabolism.

## Introduction

With over 200 million cases every year, malaria remains a major burden to global health^1^. Drug-resistant malaria parasites are increasingly common^1^, emphasizing the importance of developing antimalarials with new targets. These can be found in the apicoplast, an endosymbiotic organelle of red algal origin that is present in most apicomplexan parasites including the *Plasmodium* genus. It harbors many essential proteins with no orthologs in humans that enable development of highly specific new drugs^2^. Specifically, disease-causing asexual blood-stage parasites rely on the methylerythritol phosphate (MEP) pathway, the sulfur utilization factor (SUF) pathway, part of the coenzyme A (CoA) biosynthesis pathway, and apicoplast housekeeping^3^. A subset of asexual blood-stage parasites differentiates into male and female gametocytes, which form gametes when they are taken up by a mosquito. After fertilization, the parasites form oocysts in the mosquito midgut, which finally produce sporozoites that migrate to the salivary glands to infect a new human host. In addition to the pathways mentioned above, transmission to the mosquito host depends on apicoplast fatty acid metabolism and heme biosynthesis^4^. Although the apicoplast is an important organelle, our molecular understanding is limited due to its absence in well-studied model organisms.

One particularly poorly characterized process is the transport of metabolites into and out of the apicoplast^5^. Given that apicoplast pathways depend on many substrates that are synthesized elsewhere in the parasite, metabolite import is crucial for apicoplast functioning. Additionally, products of apicoplast metabolism that are required outside the organelle, such as isopentenyl pyrophosphate (IPP) and CoA, need to be exported^6,7^. From the four apicoplast membranes, the inner and the outer membrane are known to harbor membrane transport proteins to facilitate exchange of metabolites^8,9^. However, most of the apicoplast transporters are unidentified and uncharacterized. Only twelve (putative) transporters have been localized to the organelle in *Plasmodium spp.* to date. Of these, only three were confirmed to be essential for asexual blood-stage growth in *Plasmodium falciparum*, the malaria parasite that causes most human deaths^1,5,10^. Studying essential apicoplast proteins is challenging since their disruption results in parasite death, but this limitation can be overcome with the PfMev parasite line.^11^ This line harbors a bright apicoplast marker and an alternative metabolic pathway that produces IPP from mevalonate, facilitating the functional characterization of essential apicoplast proteins including transporters.

In this study, we characterized three members of the major facilitator superfamily in *P. falciparum*: monocarboxylate porters 1 and 2 (MCP1 and MCP2) and unidentified major facilitator (UMF). In previous work, MCP1 and UMF were suggested to be candidate vitamin B_5_ transporters on the parasite membrane, based on predicted essentiality, mRNA expression profiles, and the lack of a recognizable targeting signal or information on the substrate^12^. MCP2, on the other hand, was predicted to localize to the apicoplast based on protein proximity labeling^13^. More recently, MCP2 was found to be stabilized by atovaquone, and therefore hypothesized to be the mitochondrial atovaquone transporter, potentially besides its native function elsewhere^14^. In the related parasite *Toxoplasma gondii*, two MCP homologs and one UMF homolog localize to the apicoplast^15–18^. Here, we demonstrate that MCP1 and UMF play an essential role in apicoplast housekeeping during asexual blood-stage development, and that MCP2 has a crucial role in parasite development in the mosquito.

## Results

### Identification of three novel apicoplast proteins

To study the localization of *MCP1* (PF3D7_0210300), *MCP2* (PF3D7_0926400), and *UMF* (PF3D7_1104800), we fused the endogenous genes in-frame with the coding sequence of a C- terminal triple FLAG tag using CRISPR-Cas9 (Figure S1A). The parental parasite line was PfMev, which harbors an apicoplast-localized HA-tagged Super Folder Green (api-SFG) protein^11^. Gene-edited parasites were single-cell sorted to obtain clonal parasite lines. Correct integration and absence of parental parasites were confirmed with diagnostic PCR (Figure S2).

MCP1-FLAG and UMF-FLAG were not detectable above background by immunofluorescence microscopy (Figure 1A), nor by western blot (Figure S3). In contrast, MCP2-FLAG showed a clear signal that seemed to localize close to or colocalize with api-SFG (Figure 1A). To improve resolution, we performed ultrastructure expansion microscopy in which MCP2-FLAG and api-SFG colocalized (Figure 1B). As an alternative approach to detect MCP1, we generated a new line in which a triple HA-tag was fused in-frame to its N-terminus in the NF54 background. In this line, we detected HA signal as an elongated tubular structure in a developing schizont, and as a spherical structure in each individual merozoite of a segmented schizont (Figure 1C). The HA-MCP1 signal did generally not overlap with MitoTracker, thus strongly suggesting apicoplast localization. Despite several attempts, we were not able to generate an (N-terminal or C-terminal) HA-tagged UMF parasite line.

**Figure 1.**
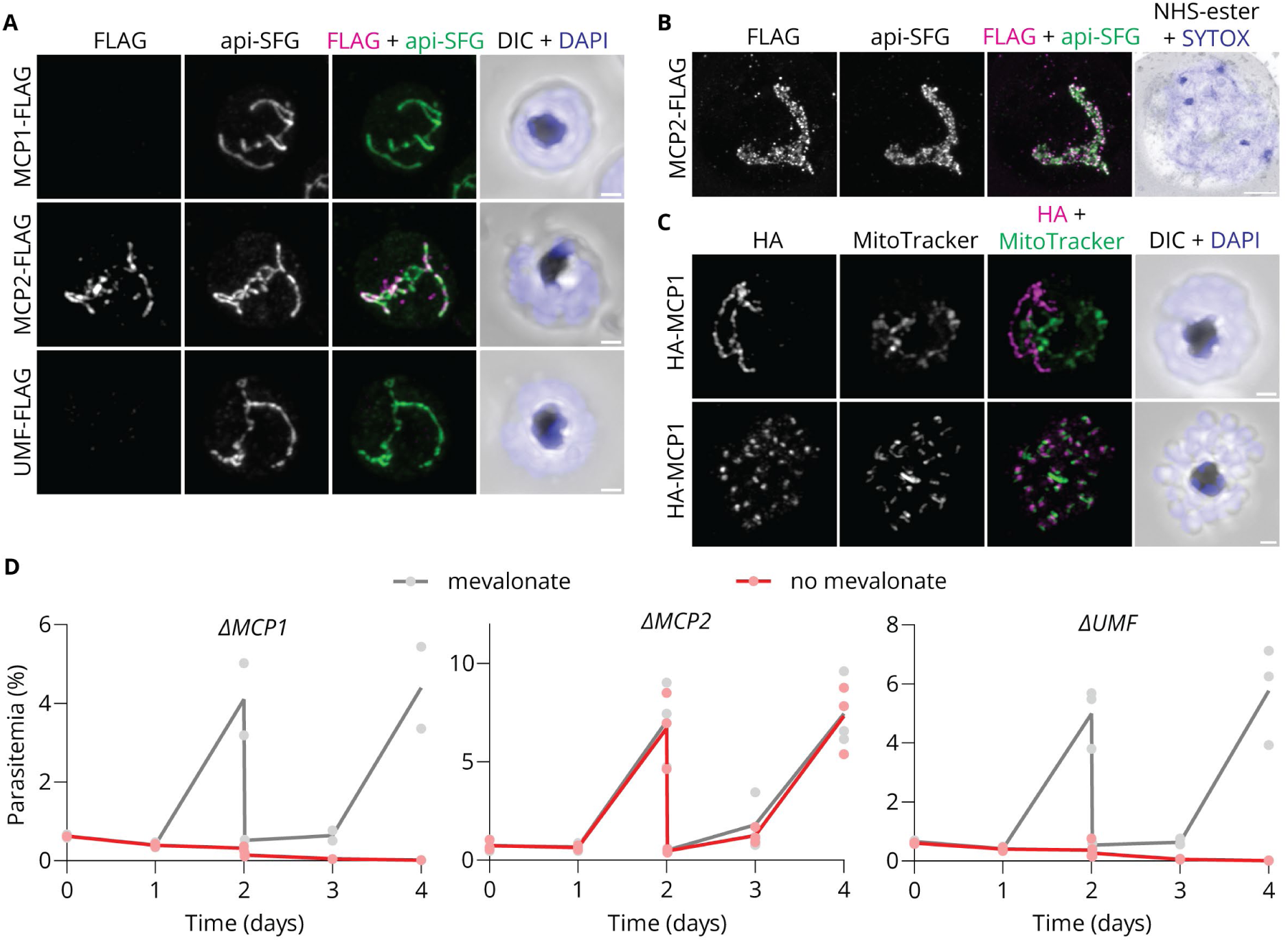
MCP1, MCP2, and UMF are apicoplast proteins. A) Immunoffuorescence microscopy of developing schizonts with C-terminal FLAG-tagged MCP1/MCP2/UMF in the PfMev background. Samples were stained with α-FLAG antibody to detect the protein of interest, α-HA antibody to visualize the SFG-HA apicoplast marker, and DAPI. Images are maximum intensity projections of Z-stacks. Brightness/contrast settings for the FLAG channel images are equal. Scale bars: 1 µm. DIC: Differential Interference Contrast. B) Ultrastructure expansion microscopy of a developing schizont with C-terminal FLAG-tagged MCP2 in the PfMev background. Samples were stained with α-FLAG antibody to detect the protein of interest, α-HA antibody to visualize the SFG-HA apicoplast marker, SYTOX, and NHS-ester. Image is a maximum intensity projection of a Z-stack. Scale bar: 4 µm. C) Immunoffuorescence microscopy of parasites with N-terminal HA-tagged MCP1 in the NF54 background. Samples were stained with α-HA antibody to detect MCP1, MitoTracker, and DAPI. Image is a maximum intensity projection of a Z-stack. Scale bars: 1 µm. D) Growth curves of synchronized asexual blood-stage PfMev parasites after deletion of a gene of interest with and without 50 µM mevalonate (gray and red, respectively). All cultures were diluted (1:20) on day 2. Each dot represents a measurement in an independent experiment; the line represents the average.

To study whether *MCP1*, *MCP2*, and *UMF* have an essential function in the apicoplast during asexual blood-stage development, we deleted the genes of interest in the PfMev background under mevalonate supplementation. Under these conditions, PfMev parasites can produce IPP with a cytosolic mevalonate pathway, bypassing the apicoplast MEP pathway and the pathways that sustain it (*i.e.* the SUF pathway and apicoplast housekeeping)^11^. We replaced all three genes with a cytoplasmic fluorescent marker gene (Figure S1) and obtained isogenic populations of *ΔMCP1*, *ΔMCP2*, and *ΔUMF* parasites using flow cytometry, which we confirmed with diagnostic PCR (Figure S2). Next, we assessed parasite growth with and without mevalonate. *ΔMCP1* and *ΔUMF* were dependent on mevalonate supplementation for growth (Figure 1D), suggesting an essential function in the apicoplast of asexual blood-stage parasites. To confirm our findings, we generated two additional parasite lines in which *MCP1* and *UMF* were disrupted by the introduction of a stop codon and a reporter gene (Figure S1). The resulting lines, *MCP1^dis^* and *UMF^dis^*, were also dependent on mevalonate for growth (Figure S4A). In contrast, *ΔMCP2* was not affected by the absence of mevalonate (Figure 1D), indicating dispensability during this stage.

### MCP1 and UMF are essential for apicoplast housekeeping

Next, we aimed to determine in which apicoplast pathway(s) MCP1 and UMF are involved. Apicoplast proteins that are essential for asexual blood-stage parasites can be involved in apicoplast housekeeping, the MEP pathway, the SUF pathway, or the CoA biosynthesis pathway. Disruptions of these pathways while supplemented with (an alternative source of) IPP can be phenotypically differentiated. A disruption in apicoplast housekeeping results in loss of both the apicoplast genome and the typical branched, tubular network morphology^2,19,20^. In contrast, blocking the MEP or SUF pathway leaves the apicoplast intact^19,20^. Disrupting CoA biosynthesis is lethal to the parasites, and unlike the other processes, this pathway cannot be bypassed with IPP supplementation^6^. We found that *ΔMCP1* and *ΔUMF* parasites supplemented with mevalonate, as substrate to produce IPP, displayed disintegration of the apicoplast network and loss of the organellar genome (Figure 2). *MCP1^dis^* and *UMF^dis^* showed the same phenotype (Figure S4B-C). These results correspond to the expected phenotype upon knockout of a gene required for apicoplast housekeeping, but do not rule out a second function in the MEP or SUF pathway.

**Figure 2.**
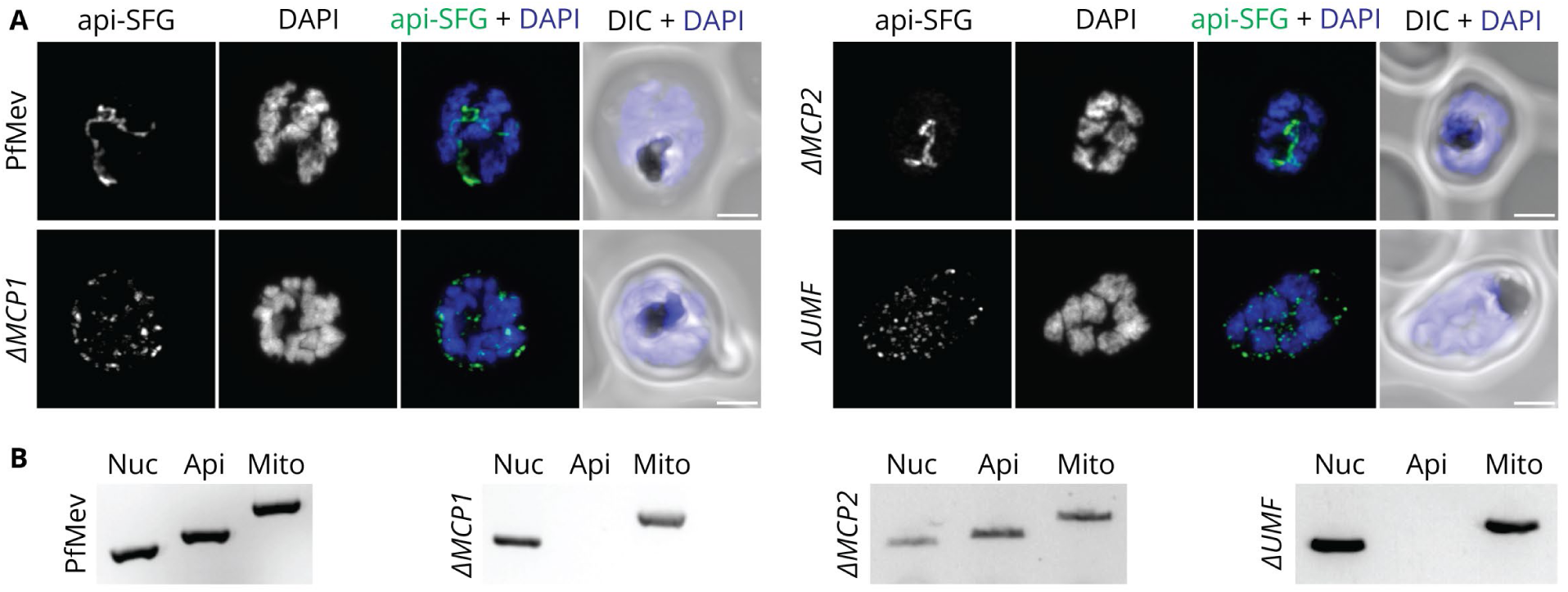
MCP1 and UMF are required for apicoplast housekeeping. A) Immunoffuorescence microscopy of developing schizonts after deletion of a gene of interest under supplementation with 50 µM mevalonate in the PfMev background to determine apicoplast integrity. Samples were stained with α-HA antibody to visualize the SFG-HA apicoplast marker and DAPI. Images are maximum intensity projections of Z-stacks. Scale bars: 2 µm. DIC: Differential Interference Contrast. B) PCR detection of nuclear (nuc), apicoplast (api) and mitochondrial (mito) genomes in parasites after deletion of a gene of interest under supplementation with 50 µM mevalonate in the PfMev background.

We selected MCP1 for further experiments using a previously validated expression vector for yeast^21^. MCP1 is a member of the monocarboxylate transporter (MCT) family, whose members generally transport small molecules such as lactate and pyruvate, although some can also transport amino acids, vitamins, and hormones^22^. In the AlphaFold 3 model of MCP1, there are three positively charged amino acid side chains pointing into the MCP1 binding pocket, suggesting a substrate with a (triple) negative charge (Figure S5A)^23^. Combined with our biological observations, we selected several candidate substrates: amino acids (variable charge, apicoplast housekeeping), isopentenyl pyrophosphate (3 negative charges, MEP pathway), pyruvate (1 negative charge, MEP pathway), and pyridoxal phosphate (2 negative charges, SUF pathway). To assess whether MCP1 transports any of these substrates, we performed a variety of transport assays based on the availability of radiolabeled substrate or deficient yeast strains (Table S1). We observed no transport activity for any of the tested substrates in the tested conditions (Figure S5).

### *MCP2* is crucial for oocyst development

Having determined that *MCP2* is dispensable for asexual blood-stage parasites, we set out to study its role in the rest of the parasite’s life cycle. Since PfMev does not form gametocytes, we generated a new *ΔMCP2* line in the NF54 background, again obtaining an isogenic population by flow cytometry (Figure S2). Twelve days after gametocyte induction, healthy-looking mature gametocytes were formed that showed no defect in the formation of exflagellation centers (Figure 3A). These gametocytes were fed to *Anopheles stephensi* mosquitoes in standard membrane feeding assays, which revealed a statistically significant 4.1-fold decrease in median oocyst intensity (NF54: 29, *ΔMCP2*: 7) while oocyst prevalence was high (>95%) for both lines (Figure 3B, S6A).

**Figure 3.**
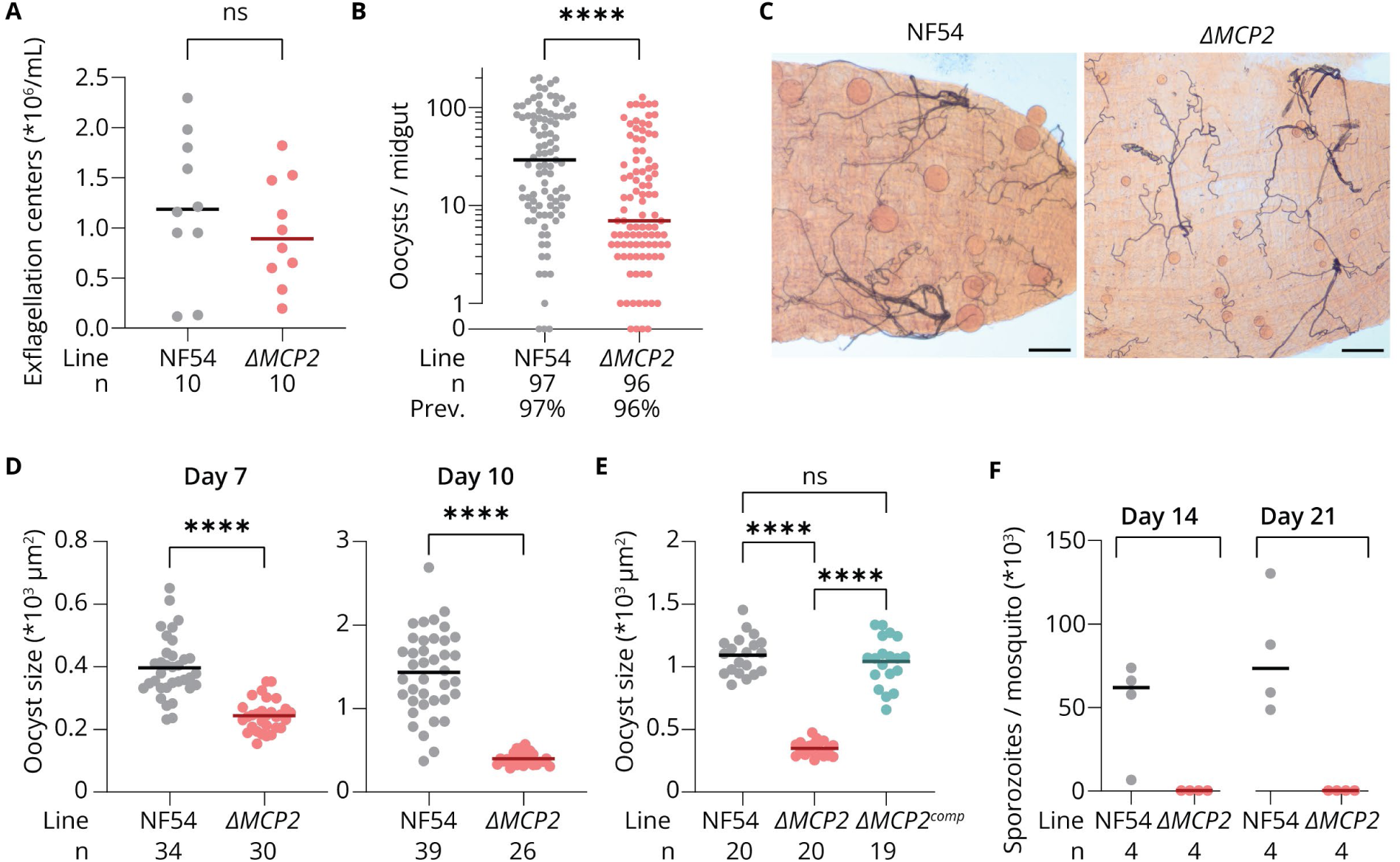
MCP2 is required for oocyst development in the mosquito. A) Million exffagellation centers per mL of gametocyte culture on day 12 after gametocyte induction of NF54 and ΔMCP2 parasites. B) Oocyst intensity in Anopheles stephensi midguts after standard membrane feeding assay with NF54 or ΔMCP2 gametocytes. Prev.: prevalence. C) Brightfield microscopy of mercurochrome-stained midguts infected with NF54 (left) or ΔMCP2 (right) oocysts at day 10 after the infectious bloodmeal. Scale bars: 0.5 mm. D) Oocyst size per mosquito at day 7 and day 10 after the infectious bloodmeal with NF54 or ΔMCP2 parasites. Each data point represents the median oocyst size in a single mosquito, mosquitoes with two or less oocysts were excluded. E) Same as D at day 10 with the additional ΔMCP2^comp^ parasite line, in which the MCP2 gene was re-inserted in the ΔMCP2 background. F) Sporozoite numbers at day 14 and day 21 after the infectious bloodmeal with NF54 or ΔMCP2 parasites. Salivary glands from 10 mosquitoes were pooled per data point to determine the average number of salivary gland sporozoites per dissected mosquito. For all graphs, the horizontal lines represent the medians (A, B, F) or the means (D, E). Statistical significance was calculated with a Mann-Whitney U test (A, B), Welch’s t test (D), or Welch’s ANOVA (E). n: number of data points, ns: not significant; ****: p < 0.0001.

*ΔMCP2* oocysts were significantly smaller on day 7 after infection (NF54: 397 µm^2^, *ΔMCP2*: 244 µm^2^), with a more pronounced difference on day 10 (NF54: 1433 µm^2^, *ΔMCP2*: 400 µm^2^) (Figure 3C-D). Between day 7 and day 10, *ΔMCP2* oocysts were still growing, increasing 56% in size compared to 283% for NF54 oocysts. To confirm that the smaller size was a result of the deletion of *MCP2*, we re-introduced the *MCP2* gene in the *ΔMCP2* line, yielding *ΔMCP2^comp^*. Reassuringly, there was no significant difference between *ΔMCP2^comp^* and NF54 oocyst size at day 10, while *ΔMCP2^comp^* oocysts were significantly larger than *ΔMCP2* oocysts (Figure 3E). In this single mosquito infection experiment, oocyst intensity also seemed to be restored in the complemented parasites (Figure S6B). Finally, we observed no *ΔMCP2* salivary gland sporozoites at day 14 and 21 after the infectious bloodmeal, indicating that oocyst maturation was not just delayed but completely blocked upon *MCP2* deletion (Figure 3F). The *ΔMCP2^comp^* parasite line did produce salivary gland sporozoites (Figure S6C).

The phenotype of *ΔMCP2* parasites closely resembles the phenotype of parasites in which *FabB/F*, a gene involved in apicoplast fatty acid metabolism, is deleted^24^. Like *ΔMCP2* parasites, *ΔFabB/F* parasites form oocysts that are smaller than wild-type oocysts and cannot produce sporozoites. Therefore, we speculated that MCP2 is involved in apicoplast fatty acid metabolism. Unfortunately, we could not replicate the previously reported expression of *Pf*MCP2 in yeast and therefore were unable to perform transport assays^21^. Instead, we attempted to rescue the *ΔMCP2* phenotype by fatty acid supplementation, taking along the previously generated *ΔFabB/F* parasites as a control^24^. We supplemented the glucose water that was supplied to the mosquitoes after the infectious bloodmeal with 250 µM BSA-coupled myristic acid and palmitic acid, which did not increase the size of *ΔFabB/F* nor *ΔMCP2* oocysts (Figure S6D). We also attempted to rescue the *ΔMCP2* phenotype with biotin supplementation, since biotin is required for fatty acid metabolism and proteins from the MCT family have been demonstrated to transport biotin^25,26^. After pre-incubation with 40 µM biotin for 24 hours, gametocytes were fed to mosquitoes in a bloodmeal containing 40 µM additional biotin, and biotin was again added to the glucose water. We observed no effect of biotin supplementation on oocyst size (Figure S6E).

Our findings indicate that MCP2 is an apicoplast protein with a crucial role in oocyst development. Surprisingly, MCP2 was recently found to be stabilized by atovaquone, prompting the hypothesis that this drug is one of its substrates^14^. To investigate this possibility, we performed drug sensitivity assays with atovaquone on NF54 and *ΔMCP2* parasites. There was no significant difference in atovaquone sensitivity between the parasites (Figure 4), indicating that atovaquone transport is unlikely to be a biologically relevant function of MCP2.

**Figure 4.**
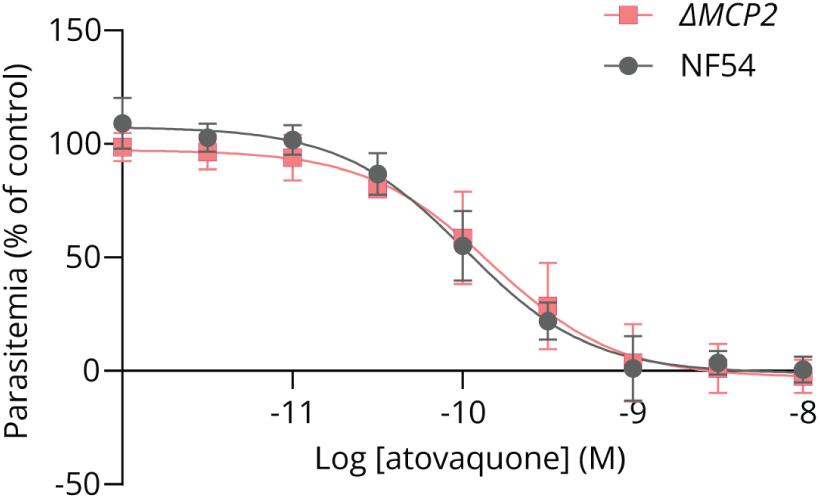
Deletion of MCP2 does not affect atovaquone sensitivity. Drug sensitivity curves of asexual blood-stage NF54 and ΔMCP2 parasites exposed to different concentrations of atovaquone for 72 hours. Averages of four independent experiments ±SD are shown. The curves were determined by nonlinear regression using a four-parameter model and the least-squares method.

### Deletion of *MCP2* affects apicoplast morphology in oocysts

Oocyst morphology can severely change upon disruption of an apicoplast protein, which can provide clues on protein function^24,27^. To visualize apicoplast morphology in *MCP2*-deficient oocysts, we generated a new parasite line, *ΔMCP2^api-SFG^*, in which the *MCP2* coding sequence was replaced for a gene encoding the apicoplast marker api-SFG (Figure S1-2). As a reference, we used the PfMev12 parasite line that we recently described, which expresses api-SFG from a silent intergenic locus^28^. At day 3 after infection, apicoplast morphology was comparable between PfMev12 and *ΔMCP2^api-SFG^* parasites (Figure 5). In both lines, we occasionally observed DNA staining colocalizing with the api-SFG signal, suggesting high levels of apicoplast DNA replication at these sites (Figure S7). At day 5, the apicoplast in PfMev12 parasites started to expand throughout the cell with a ‘beads-on-a-string’ morphology. The *ΔMCP2^api-SFG^* apicoplast remained smaller in general, and the ‘beads’ also appeared smaller. By day 7, the apicoplast in PfMev12 parasites formed an extensive tubular network throughout the oocyst, while the *ΔMCP2^api-SFG^* apicoplast remained small with short, thin extensions. At day 10, some PfMev12 oocysts presented an extremely extensive and intricate network, while in other oocysts, the apicoplast had already segmented to produce small elongated apicoplasts for each individual sporozoite. At this timepoint, the *ΔMCP2^api-SFG^* apicoplast remained relatively small and sometimes fragmented. These data show that MCP2 is required for apicoplast development in the oocyst, and is in line with the hypothesis that MCP2 facilitates apicoplast fatty acid metabolism.

**Figure 5.**
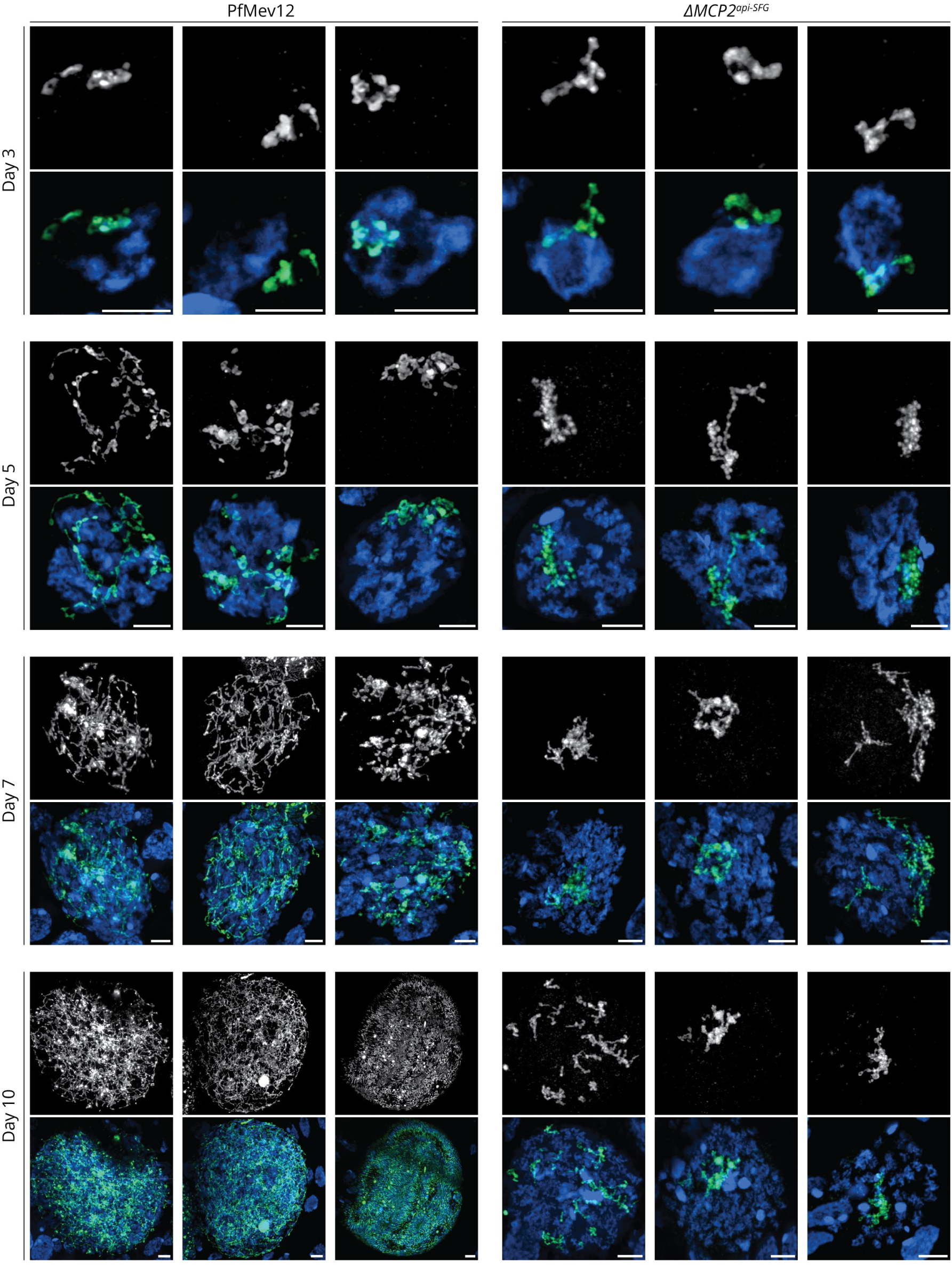
Apicoplast morphology in oocysts is severely affected upon deletion of MCP2. Fluorescence microscopy of oocysts within dissected mosquito midguts, 3, 5, 7, and 10 days after an infectious bloodmeal with PfMev12 or ΔMCP2^api-SFG^ parasites. Grayscale images show api-SFG signal. Composite images show api-SFG signal in green and Hoechst signal in blue. Images are maximum intensity projections of Z-stacks. Scale bars: 4 µm.

## Discussion

The *Plasmodium* apicoplast depends on transporters to import and export metabolites but the apicoplast transportome is very poorly characterized^5^. Here, we report the identification of three new apicoplast proteins: MCP1, MCP2, and UMF, all members of the major facilitator superfamily, bringing the total number of (putative) transporters with confirmed apicoplast localization in malaria parasites to fifteen^5,10^.

*MCP1* and *UMF* are essential during asexual blood-stage development and their deletion results in aberrant apicoplast morphology and loss of the organellar genome. These results indicate a role in apicoplast housekeeping, which encompasses organelle biogenesis pathways such as DNA replication, transcription, and translation, and organelle maintenance processes such as ion homeostasis and redox balance. Consequently, MCP1 and UMF may transport substrates used in these processes including nucleotides, amino acids, s-adenosyl methionine, and (derivatives of) vitamin B_2_, B_6_, and B_9_^5^. To complicate matters further, the observed phenotypes of a disintegrated apicoplast and loss of its genome upon knockout do not rule out a second function of MCP1 and/or UMF in another apicoplast pathway. MCP1 and/or UMF may thus have two functions (or more), for example import of a vitamin for housekeeping and export of IPP from the MEP pathway. Similar transporters also regularly have more than one substrate^22^. Making use of yeast expressing *Pf*MCP1, we performed a variety of transport assays for several candidate substrates to no avail. Also untargeted metabolomics did not lead to the identification of a substrate (data not shown). It remains unclear whether the relevant substrate(s) was/were not included in the assays, or whether the yeast-expressed *Pf*MCP1 is inactive, for example because it is incorrectly folded, lacks critical post-translational modifications, or requires a cofactor unavailable in the system used.

*MCP2* is essential for parasite transmission and its deletion results in a block of oocyst development. Currently, only two apicoplast pathways are known to be critical for transmission and not for asexual blood-stage development. Heme biosynthesis is required for male gamete formation, which was unaffected in *ΔMCP2* parasites, ruling out an essential role in this process. In contrast, apicoplast fatty acid metabolism is required for oocyst development and sporozoite formation^24^. Deletion of *FabB/F*, which disrupts apicoplast fatty acid metabolism, results in smaller oocysts with abnormal apicoplast morphology that do not produce sporozoites. Since *ΔMCP2* parasites display a similar phenotype, it seems plausible that MCP2 is involved in fatty acid metabolism, possibly through transport of candidate substrates such as biotin and/or TPP^5^. Unfortunately, we were not able to confirm this hypothesis by supplementation of fatty acids or biotin to the mosquitoes. Perhaps the supplemented compounds did not reach their target, for example due to insufficient uptake from the sugar meal or lack of scavenging by the oocysts^29^. There are also indications that MCP2 is not (only) involved in apicoplast fatty acid metabolism. For example, *ΔMCP2* oocysts did not exactly phenocopy *ΔFabB/F* oocysts, which appear slightly bigger, although this was not statistically significant. Furthermore, *ΔMCP2 P. berghei* parasites display severely impaired sporozoite formation (Huppertz, PhD thesis, 2025), while apicoplast fatty acid metabolism is dispensable for *P. berghei* development in *An. stephensi* mosquitoes^30^. Therefore, in addition to its potential function in fatty acid metabolism, MCP2 might also play a role in one or more other apicoplast pathways.

Other Apicomplexa harbor orthologs of *MCP1*, *MCP2*, and *UMF* that can potentially aid in finding the substrates for the *P. falciparum* transporters. The *T. gondii* genome contains a clear ortholog of UMF, TGME49_309580, which is confirmed to localize to the apicoplast but its substrate is unknown^16^. There is more data on ortholog group OG7_0006513 (OrthoMCL DB 7.0), of which *MCP1* and *MCP2* are the only *P. falciparum* members. *T. gondii* has three members: TGME49_233540 (*AMT1*), TGME49_297245 (*AMT2*), and TGME49_253130. AMT1 and AMT2 are confirmed to localize to the apicoplast, while TGME49_253130 localizes elsewhere^15–17^. The functions of *TgAMT1* and *TgAMT2* are the subject of controversy. One study reported that they form a complex that transports pyruvate, but another observed distinct phenotypes upon depletion, suggesting that they have different substrates.^15,17^ It seems unlikely that the MCPs form a pyruvate-transporting complex in *P. falciparum,* since we found distinct phenotypes upon deletion of the MCPs. Furthermore, pyruvate import is possibly redundant in the *Plasmodium* apicoplast^5,19^ and the MCPs do not comigrate in native protein gel electrophoresis^31^. Unlike *T. gondii*, *Cryptosporidium parvum* has lost its apicoplast but remarkably still harbors two MCP orthologs, cgd4_380 and cgd7_2450. Coincidingly, *Cryptosporidium* has lost several biosynthesis pathways, instead scavenging essential metabolites from its host. Therefore, the required transporters on the *Cryptosporidium* cell membrane are expected to functionally partially overlap with the transporters on the *Plasmodium* apicoplast membrane. If the MCP functions would be conserved between *Cryptosporidium* and *Plasmodium*, their substrate may be a compound that *Cryptosporidium* scavenges from the host, while the *Plasmodium* apicoplast exchanges it with the parasite cytoplasm. Compounds that meet these criteria are IPP, flavin adenine dinucleotide (FAD), pyridoxal-5’-phosphate (PLP), and thiamine pyrophosphate (TPP)^32,33^. FAD and PLP are required for protein translation in the apicoplast by facilitating tRNA modifications, and could therefore be prioritized as candidate substrates for MCP1^34^. TPP is a cofactor of 1-deoxy-D-xylulose 5-phosphate synthase, the first enzyme of the MEP pathway of which the final, exported product is IPP, so TPP and IPP are also candidate substrates for MCP1^35^.

MCP1, MCP2, and UMF are members of a growing group of proteins that are confirmed to localize to the apicoplast but lack the canonical apicoplast-targeting signal^36^. These proteins do not undergo the typical N-terminal processing^8,10^, which is in line with our observation that N-terminal tagging of MCP1 yields apicoplast-localized signal. Apicoplast proteins without a signal peptide are hypothesized to localize to the outer apicoplast membrane^37^, which was demonstrated for the *P. falciparum* outer triose phosphate transporter (*Pf*oTPT)^8^. Correct localization of *Pf*oTPT depends on a tyrosine residue in the cytoplasmic N-terminus of the protein and some unknown factors^37^. Indeed, MCP1, MCP2, and UMF also have tyrosine residues at positions 13, 11, and 28, respectively, before the first predicted transmembrane domain. If MCP1, MCP2, and UMF indeed specifically localize to the outer apicoplast membrane, it remains unclear how their substrates pass the inner apicoplast membranes.

In this study, we identified three novel apicoplast transporters, and show that PfMev12 is a transmissible parasite line that can serve as a valuable tool to study apicoplast proteins that are essential in the mosquito. Taken together, our work marks an important step towards the full characterization of the *Plasmodium* apicoplast transportome. From a translational perspective, our results indicate that MCP1 and UMF might be interesting targets for novel antimalarials, while MCP2 could be an attractive target for transmission-blocking interventions.

## Materials and Methods

### Molecular cloning

All kits and enzymes were used according to manufacturer’s instructions. All PCRs were performed using PrimeSTAR GXL DNA Polymerase (Takara Bio) and primers from Sigma-Aldrich or Biolegio (Table S2). Genomic DNA (gDNA) was obtained using the ǪIAamp DNA Blood Mini Kit (Ǫiagen). All restriction enzymes were purchased from NEB. Sequences were confirmed by Sanger sequencing (Baseclear) or Plasmidsaurus whole-plasmid sequencing (Oxford Nanopore, R10.4.1).

The homology-directed repair (HDR) templates were generated by traditional restriction enzyme digestion and ligation and/or Golden Gate Assembly (GGA) using the NEBridge Golden Gate Assembly Kit (BsmBI-v2) (Table S3). The starting plasmids were pRF0506 (previously generated by removing all type IIS restriction enzyme recognition sites from pUC19 by site-directed mutagenesis^38^), pRF0205 (generated by replacing the gene for mitochondrial mScarlet with a gene for cytoplasmic mScarlet under the PfH2B promoter in plasmid pRF0139 as previously described^39^), pRF0508 (previously generated by combining the PfH2B promoter, the mScarlet coding sequence, and a bidirectional UTR in a GGA reaction^38^), pRF0674 (generated in the same way as pRF0508 with a *Pf*U60s promoter instead of a *Pf*H2B promoter), and pGGAselect (NEB).

For the Cas9 guide plasmids, two complementary oligonucleotides encoding the guides were phosphorylated with T4 polynucleotide kinase (NEB), annealed, and ligated into BbsI-digested pRF0277^28^, a plasmid derived from pMLB626 (kind gift from Marcus Lee)^40^ with an additional negative selection marker cassette (*Sc*FCY1/*Sc*FUR1)^41^.

### Parasite culture and transfections

*P. falciparum* cultures were maintained in 5% O+ human red blood cells (Sanquin) in RPMI1640 medium with 25 mM HEPES, 25 mM NaHCO_3_, 333 µM hypoxanthine, and 10% human type A serum (Sanquin) (complete medium) at 37 °C with 3% O_2_ and 4% CO_2_^42^. Parasites were synchronized using sorbitol if required for transfections or gametocyte induction^43^.

Transfections were performed on ring-stage parasites as previously described^44^. In short, 60 µg linearized HDR template and one or two guide plasmids (60 µg per plasmid) (Figure S1) were ethanol precipitated, dissolved in cytomix (10 mM K_2_HPO_4_/KH_2_PO_4_ pH 7.6, 120 mM KCl, 0.15 mM CaCl_2_, 5 mM MgCl_2_, 25 mM HEPES, 2 mM EDTA), and added to a 5% ring-stage parasite culture. After electroporation (310 V, 950 µF), the parasites were allowed to recover for 4 h before selection was started with 2.5 nM WR99210 (Jacobus Pharmaceuticals) for 5 days^45^. For the gene deletion and disruption lines in the PfMev background, 50 µM mevalonate (supplied as 50 µM mevalonolactone (Sigma-Aldrich)) was added to the medium from the start of drug selection onwards. To facilitate re-transfection to reintroduce the *mcp2* gene in the *ΔMCP2* line, the parasites were treated with 1 µM 5-fluorocytosine (Sigma-Aldrich) for 5 days to remove any remaining guide plasmid before the second transfection^41^.

To obtain clonal populations of the FLAG-tagged and HA-tagged lines, schizonts were stained with 1 µg/mL Rhodamine123, after which single cells were sorted into a round-bottom 96-well plate containing 100 µL complete medium and 3% hematocrit per well using a FACSAria III Cell Sorter (BD Biosciences) or a Cytoflex SRT Benchtop Cell Sorter (Beckman Coulter). Complete medium was replaced every three days until parasites were detectable by blood smear microscopy, after which clonal parasite cultures were scaled up. To obtain isogenic populations of the gene deletion and disruption lines, 250 parasites positive for the fluorescent mScarlet or api-SFG marker were sorted with one of the abovementioned cell sorters. Successful integration and absence of parental line was confirmed with PCR.

### Gametocyte culturing and mosquito infections

To induce gametocytogenesis, trophozoite-stage parasite cultures at 1% parasitemia were treated with 0.5% AlbuMAX-II (Gibco) in RPMI1640 with 25 mM HEPES, 25 mM NaHCO_3_, and 100 mM hypoxanthine for 36 h as previously described^46^. After induction, gametocytes were cultured in complete medium in a semi-automatic shaker set-up^47^. Exflagellation was quantified on day 12 after induction by mixing the culture 1:1 with 200 µM xanthurenic acid in complete medium, followed by 15 min incubation at RT in a humid chamber before counting exflagellation centers (aggregates of 3 or more red blood cells) using an improved Neubauer chamber.

*An. stephensi* mosquitoes (Nijmegen Sind-Kasur strain^48^) were reared at 30 °C and 80% relative humidity under a 12 h:12 h light-dark cycle with *ad libitum* access to 5% glucose water. Standard membrane feeding assays were performed as described previously^49^. In short, equal amounts of mature gametocytes were added to a bloodmeal. Unfed and partially fed mosquitoes were removed, and blood-fed mosquitoes were maintained at 26 °C. To assess oocyst prevalence, intensity and size, midguts were dissected on day 7 and 10 after infection and stained with 1% mercurochrome for 10 min, followed by imaging with a ZEISS Axio Scope.A1 microscope and a Basler acA5472-17uc camera. Images were analyzed using a previously published image pipeline^50^ in combination with a script to automate image mapping (https://gist.github.com/gregorysabatino/89e3d7252b0358058110ef97f89bead0/). For figure 3D, oocysts were manually measured using Fiji^51^. For figures 3B, 3E, and S6, oocysts were analyzed using the Oocystmeter^50^, followed by thorough manual curation to add missed oocysts and remove wrong annotations. To quantify salivary gland sporozoites, the salivary glands of 10 mosquitoes were dissected and pooled in PBS, grinded, and loaded onto an improved Neubauer chamber. The sporozoites were left for 20 minutes to settle and then counted.

For the fatty acid supplementation, myristic acid (C14:0, Carl Roth) and palmitic acid (C16:0, Carl Roth) were coupled to fatty acid-free BSA Fraction V (Carl Roth) by sonication, and then diluted to a final concentration of 250 µM in 5% glucose water, which was supplied to the mosquitoes from one day after the bloodmeal onwards. For biotin supplementation, 40 µM biotin (Sigma Aldrich) was supplemented to the gametocytes one day before the bloodmeal, 40 µM biotin was added to the bloodmeal, and 40 µM was added to the glucose water, freshly made each day and supplied from one day after the bloodmeal onwards.

### Fluorescence microscopy

Primary antibodies used were mouse α-FLAG M2 (F1804, Sigma-Aldrich, 1:500 for immunofluorescence assay (IFA), 1:250 for ultrastructure expansion microscopy (U-ExM)) and rat α-HA 3F10 (11867423001, Roche, 1:500 for IFA, 1:250 for U-ExM). Secondary antibodies used were chicken-α-rat Alexa Fluor 488 (Invitrogen A-21470, 1:1000 for IFA, 1:500 for U-ExM) and goat- α-mouse Alexa Fluor 568 (Invitrogen A-11031, 1:1000 for IFA, 1:500 for U-ExM). Fluorescence microscopy was performed on a ZEISS LSM900 confocal microscope with Airyscan using a 63x oil objective and 405, 488, 561, and 633 nm laser excitation. Airyscan processing was done with ZEISS ZEN Blue. Further analysis was performed in Fiji^51^.

For IFA, cells were stained with 100 nM MitoTracker Orange CMTMRos (Thermo Fisher) in complete medium for 30 min at 37 °C if visualization of the mitochondrion was required. Then, they were settled onto poly-L-lysin-coated coverslips (Corning), fixed with 4% methanol-free formaldehyde (Thermo Fisher) and 0.0075% EM-grade glutaraldehyde (Thermo Fisher) in PBS, permeabilized with 0.1% Triton X-100 for 10 min, blocked with 3% BSA in PBS for 1 h, stained with primary and secondary antibodies for 1 h with three PBS washes in between, stained with 1 µM DAPI (Thermo Fisher) for 10 min, and mounted on slides using Fluoromount G (Thermo Fisher).

For U-ExM, a previously published protocol was followed with minor modifications^52^. Briefly, late asexual blood-stage parasites were enriched using a MACS LS column (Miltenyi Biotec).^53^ Then, they were fixed as described above, incubated in anchoring solution (1.2% acrylamide (Sigma-Aldrich), 2% formaldehyde (Sigma-Aldrich)) at 37 °C overnight, followed by gelation in 19% sodium acrylate (AK Scientific), 10% acrylamide, 0.1% N,N’-methylenebisacrylamide (Sigma-Aldrich), 0.5% tetramethylethylenediamine (Sigma-Aldrich) and 0.5% ammonium persulfate (Sigma-Aldrich). Gels were incubated in denaturation buffer (200 mM SDS, 200 mM NaCl, 50 mM Tris, pH 9) for 90 min at 95 °C and then stained with antibodies, DyLight 405 NHS Ester (Thermo Fisher, 1:250) and SYTOX Deep Red (Thermo Fisher, 1:1000). Prior to imaging, gels were expanded by incubation in milli-Ǫ water.

For fluorescence microscopy of oocysts, mosquito midguts were fixed with 4% paraformaldehyde for 1 h at RT, washed with PBS, stained with 10 µg/mL Hoechst 33342 (Thermo Fisher) for 1 h at RT, washed with PBS, mounted on glass slides in Fluoromount G, and imaged on the same day.

### Western blotting

Mixed asexual blood-stage parasites were released from red blood cells by treatment with 0.06% saponin (Thermo Fisher) in PBS and washed with PBS. Parasites were then lysed in 1x NuPAGE LDS buffer (GenScript) and incubated at 37 °C for 30 min. Samples were run on a 10% bis-Tris SDS-Page gel (SurePAGE) with Precision Plus Dual Color Standards (Bio-Rad) as a reference. Following electrophoresis, proteins were transferred onto a 0.45 µm PVDF membrane (Bio-Rad) with the Trans-Blot Turbo transfer system (Bio-Rad). The membrane was blocked with 3% BSA in PBS, followed by incubation with primary antibody in 1% BSA in PBS with 0.05% Tween-20 at 4 °C overnight. The membrane was then washed, incubated in secondary antibody at RT for 1 h, washed, and developed with Clarity Max ECL substrate (Bio-Rad) to image on an ImageǪuant LAS 4000 (GE Healthcare). Primary antibodies used were mouse α-FLAG M2 (F1804, Sigma-Aldrich, 1:1000) and rabbit α-*Pf*HSP70 (SPC-186, StressMarq, 1:1000). Secondary antibodies used were HRP-conjugated rabbit-α-mouse (Dako, 1:2000) or goat-α-rabbit (Dako, 1:2000).

### Growth assays

Asexual blood-stage parasite growth was measured with a flow cytometry-based growth assay. Cultures were sorbitol-synchronized twice, with 16 h in between, to obtain tightly synchronized young ring-stage parasites. The assay cultures were then set up at 0.5% parasitemia with or without 50 µM mevalonate (supplied as 50 µM mevalonolactone) supplemented to the medium. Two days after setup, all cultures were diluted (1:20 ratio) with fresh complete medium and RBCs. Each day of the assay, samples were taken, fixed in 0.25% glutaraldehyde (Thermo Fisher) in PBS, and stored at 4 °C up to two weeks before measurements. Parasitemia in all samples was measured on the same day, by staining the DNA with SYBR Green (Thermo Fisher, 1:10,000 diluted) for 30 min at 37 °C, followed by quantification of the infected cells with flow cytometry on a CytoFLEX 13-color flow cytometer. Data was analyzed using FlowJo v10.

### Detection of nuclear/mitochondrial/apicoplast genome

To investigate the presence of the apicoplast organellar genome after gene deletion or disruption, part of the apicoplast-encoded *sufB* gene (PF3D7_API04700) was amplified by PCR using previously published primers (Table S2)^11^. As positive controls, part of the nuclear-encoded *ldh* gene (PF3D7_1324900) and the mitochondrion-encoded *cox1* gene (PF3D7_MIT02100) were amplified.

### Drug sensitivity measurements

Drug sensitivity was measured in a replication assay as previously described^54^. In short, parasites were incubated with different concentrations of atovaquone in a 96-well plate for 72 h. Parasites were then lysed in presence of SYBR Green, followed by fluorescence intensity measurements using a BioTek Synergy 2 plate reader. Values were normalized to a 1 µM dihydroartemisinin control (0% growth) and a 0.1% DMSO control (100% growth).

### Protein modelling

The MCP1 structure was predicted by AlphaFold 3^23^ and analyzed with ChimeraX^55^.

### Transport assays in yeast

To measure transport of IPP, pyruvate, and amino acids, the yeast strain used was *Saccharomyces cerevisiae* W303-1A Δ*jen1* Δ*ady2* (MATa, *can1-100 ade2-1 his3-11,15 leu2-3,112 trp1-1 ura3-1 jen1::kanMX4 ady2::hphMX4*) kindly provided by M. Casal^56^. Yeast was transformed by the LiAc/ssDNA/PEG method^57^ with a previously published pDR196-based expression construct, which was confirmed to facilitate expression of *Pf*MCP1 on the yeast plasma membrane^21^. Empty pDR196 plasmid was used as a negative control. Transformed yeast was grown at 29 °C, 220 rpm in selective synthetic defined medium with Difco Yeast Nitrogen Base (Becton Dickinson), 0.5% (w/v) ammonium sulfate, 2% (w/v) glucose, 20 mg/L L-histidine, 10 mg/L L-tryptophan, 100 mg/L L-leucine, and 25 mg/L adenine.

Direct transport assays using radiolabeled IPP and pyruvate were performed as previously described^21^. In short, yeast was exposed to a substrate gradient (0.5 mM for IPP, 1 mM for pyruvate) spiked with 0.04 µCi ^14^C-labeled substrate (Hartmann Analytic) for a defined period of time. The substrate solution was then removed by filtration, the yeast was washed and radioactivity was assessed as a measure of substrate uptake.

Transport of amino acids was tested by detecting transport-induced changes in yeast cell volume using stopped-flow light scattering as previously described^58^. In short, yeast was exposed to a 300 mM substrate gradient within a stopped-flow mixing system (SFM-2000, BioLogic), inducing a quick efflux of water from the yeast cells, detected as an increase in light scattering. Transport of amino acids would then be detected as a gradual decrease in light scattering after the initial peak, reflecting the influx of water along with the amino acid.

To assess transport of pyridoxine (PN) and PLP, the yeast strain used was *S. cerevisiae* W303-1A Δ*snz1* Δ*sno1* Δ*tpn1* (MATa, *can1-100 ade2-1 his3-11,15 leu2-3,112 trp1-1 ura3-1 snz1- sno1::Sphis5 tpn1::kanMX4*) kindly provided by Jürgen Stolz^59^. This strain lacks crucial components of the vitamin B6 salvage and uptake pathways and hardly grows at concentrations of 0.2 µM PN or lower. The yeast was transformed as described above and grown in ‘home-made’ synthetic defined medium with the same components described above except for PN, which was added to the medium in the desired concentration at the setup of the experiment.

## Acknowledgements

We would like to thank Sean Prigge for sharing the PfMev parasite line. We appreciate the help of Rianne Stoter, Geert-Jan van Gemert, Laura Pelser-Posthumus, Jolanda Klaassen, Astrid Pouwelsen, and Freia-Raphaella Lorenz with the mosquito rearing, infections, and dissections. We thank Mathieu Brochet and Ravish Rashpa for their lessons in ultrastructure expansion microscopy. We highly appreciate the efforts of Robert Jansen and Pingping Zhu in using untargeted metabolomics to identify the transporter substrates. We thank Sofia Tsalkitzidou for her assistance. We are grateful for the insightful discussions about the project with Dominique Soldati-Favre, Joachim Kloehn, and Szilamér Koszti. We also acknowledge the Radboudumc Technology Centers for Flow Cytometry and Microscopy for the use of their machines.

## Funding Statement

This work was supported by the Radboud University Medical Center (Radboudumc Individual PhD for Masters round 2021 to LJA, Radboudumc Individual PhD for Masters round 2021 to ETB), the European Molecular Biology Organization (Scientific Exchange Grant to LJA), Radboud University (Christine Mohrmann Stipend to LJA), the Dutch Research Council (grant ID: https://doi.org/10.61686/HHGJT46263 to TWAK; grant ID https://doi.org/10.61686/QCMAK98128 to LEdV), the Deutsche Forschungsgemeinschaft (Be2253/8 to EB), and the Dr. Hilmer Stiftung (T0146 – 42.804 to SB). Ultrastructure expansion microscopy was funded by a Radboud Community for Infectious Diseases grant (RCI Grant 2024) awarded to TWAK.

## Notes

### Competing Interest Statement

The authors have declared no competing interest.

