## Supplemental Info for "Identification of three apicoplast transporters with crucial roles in the malaria parasite life cycle"

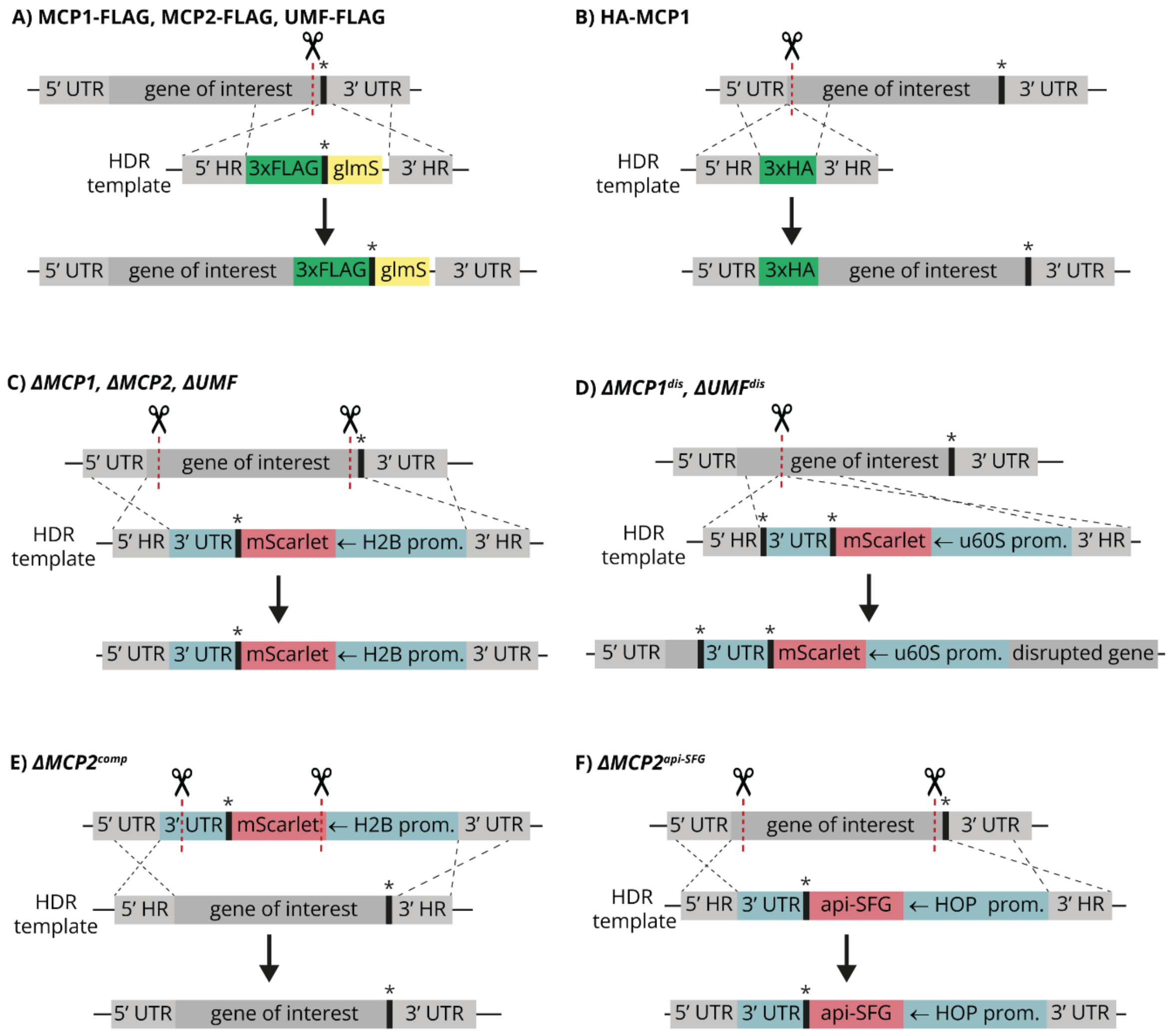


***Figure S1 – Design of genetically modified parasite lines to study* MCP1, MCP2, *and* UMF.** *A) C-terminal FLAG-tag design. B) N-terminal HA-tag design. C) Gene deletion design. D) Gene disruption design.* MCP1 *was disrupted 249 bp from the start codon.* UMF *was disrupted 150 bp from the start codon. E) Gene complementation design. F) Gene deletion with apicoplast marker design. Scissors indicate CRISPR-Cas9 guide target sites. Asterisks indicate stop codons. HDR: Homology-Directed Repair, HR: Homology Region, UTR: untranslated region, H2B prom.: promoter region of histone H2B (PF3D7_1105100), u60S prom.: promoter region of ubiquitin-60S ribosomal protein L40 (PF3D7_1365900), HOP prom.: promoter region of Hsp70/Hsp90 organizing protein (PF3D7_1434300), api-SFG: HA-tagged Super Folder Green with N-terminal apicoplast localization signal.*

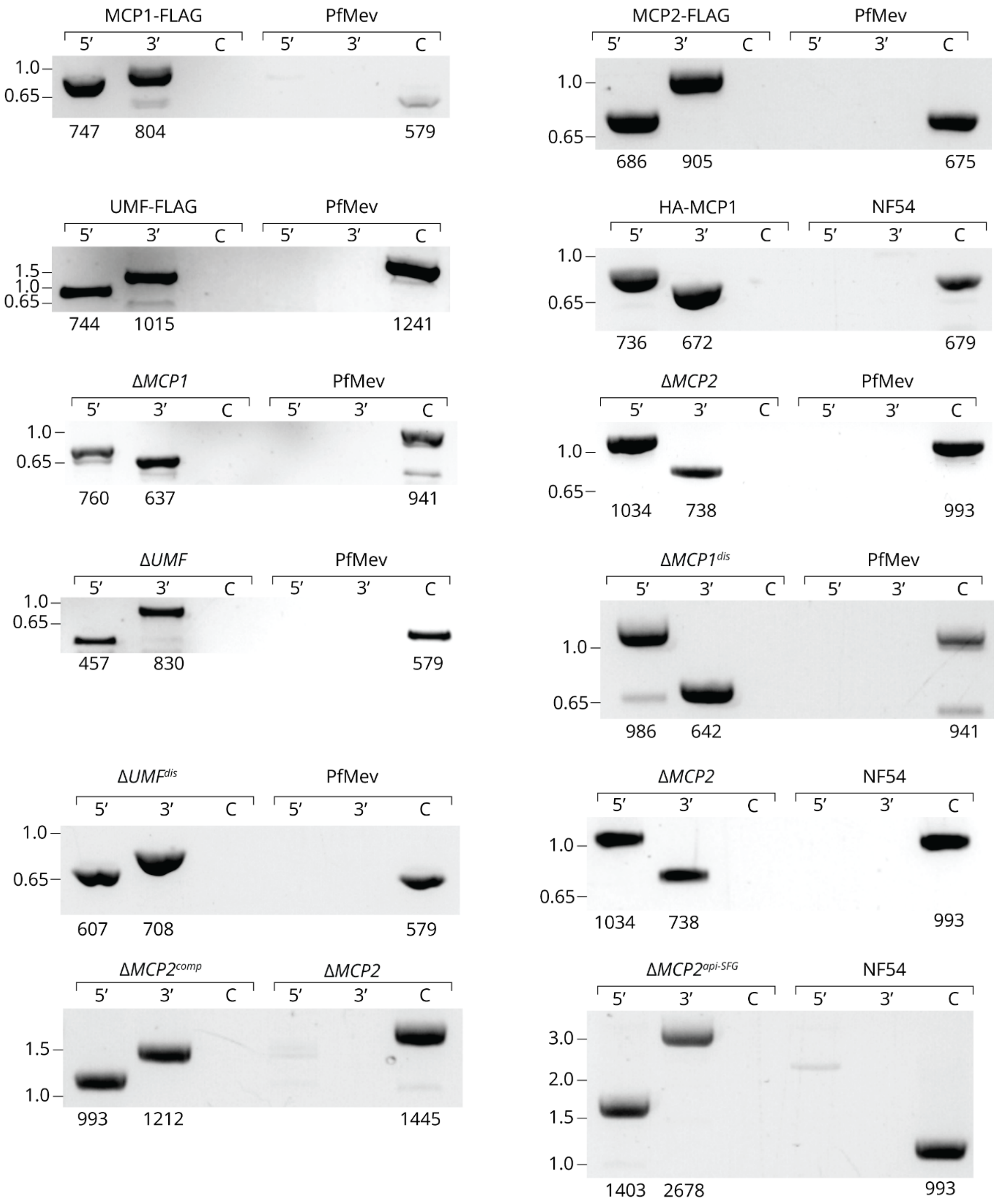


***Figure S2 – Diagnostic PCRs of genetically modified parasite lines.*** *For each line, PCR reactions were performed to confirm 5' and 3' integration of the homology-directed repair construct and a control (C) reaction to check the presence of parental parasites. The same reactions were performed on the parental parasites as a control. The approximate sizes (in kb), based on Thermo Fisher 1 Kb Plus DNA Ladder, are indicated on the left of the images. The expected product sizes (in bp) are indicated below the images.*

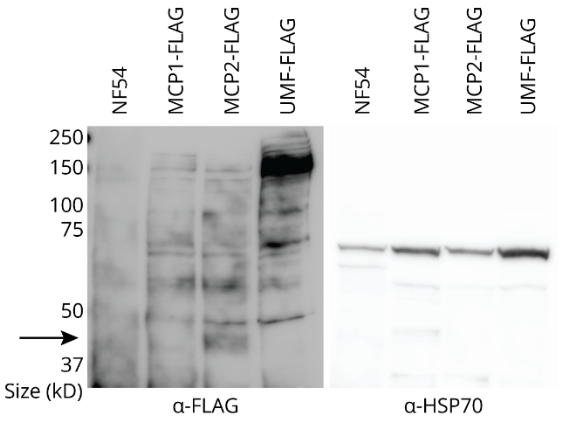


***Figure S3 – Western blot of FLAG-tagged proteins of interest.*** *Mixed asexual blood-stage parasite lysates were run on a non-reduced SDS-PAGE gel and western blots were stained with α-FLAG antibody to detect the tagged protein of interest or α-HSP70 antibody as a loading reference. MCP2-FLAG (predicted size 61 kDa) seems to appear as a faint band around 45 kDa (indicated by the arrow). MCP1-FLAG and UMF-FLAG (predicted sizes 52 kDa and 49 kDa, respectively) cannot be discerned from the background bands.*

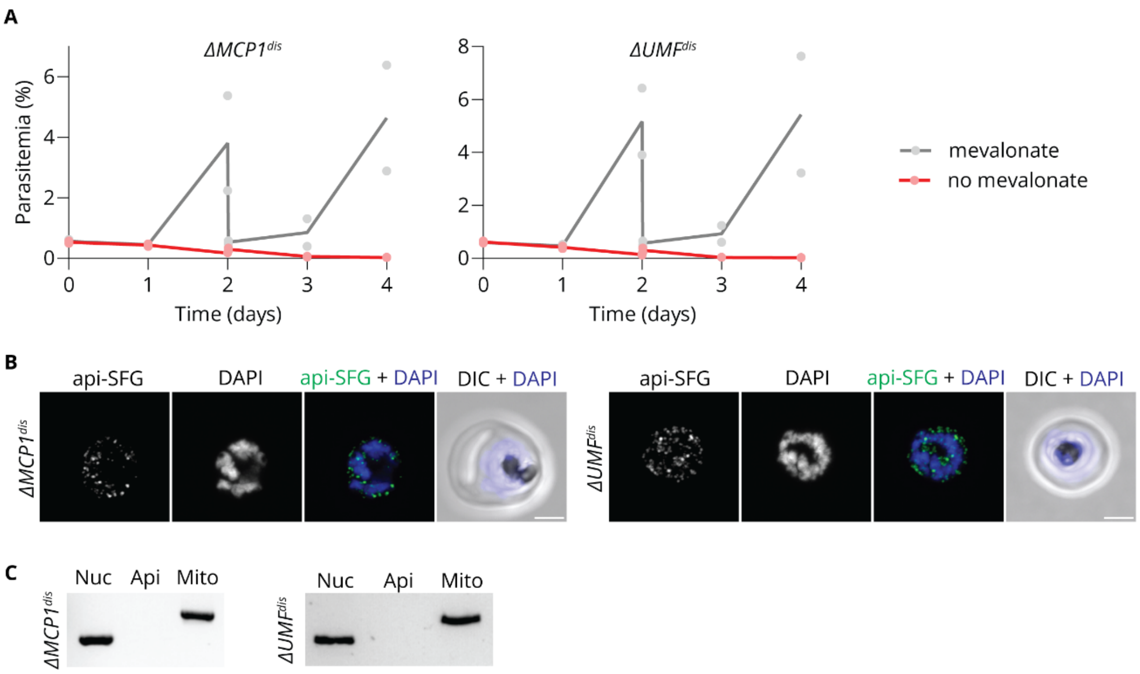


***Figure S4 – Disruption of* MCP1 *and* UMF *disturbs apicoplast housekeeping.*** *A) Growth curves of synchronized asexual blood-stage PfMev parasites after disruption of* MCP1 *or* UMF *cultured with or without 50 μM mevalonate (gray and red, respectively). All cultures were diluted (1:20) on day 2. Each dot represents a measurement in an independent experiment; the line represents the average. B) Immunofluorescence microscopy to determine apicoplast integrity of developing*

*schizonts after disruption of* MCP1 *or* UMF *in the PfMev background under supplementation with 50 µM mevalonate. Samples were stained with α-HA antibody to visualize the SFG-HA apicoplast marker and DAPI. Images are maximum intensity projections of Z-stacks. Scale bars: 2 µm. DIC: Differential Interference Contrast. C) PCR detection of nuclear (nuc), apicoplast (api) and mitochondrial (mito) genomes in parasites after disruption of* MCP1 *or* UMF *under supplementation with 50 µM mevalonate in the PfMev background.*

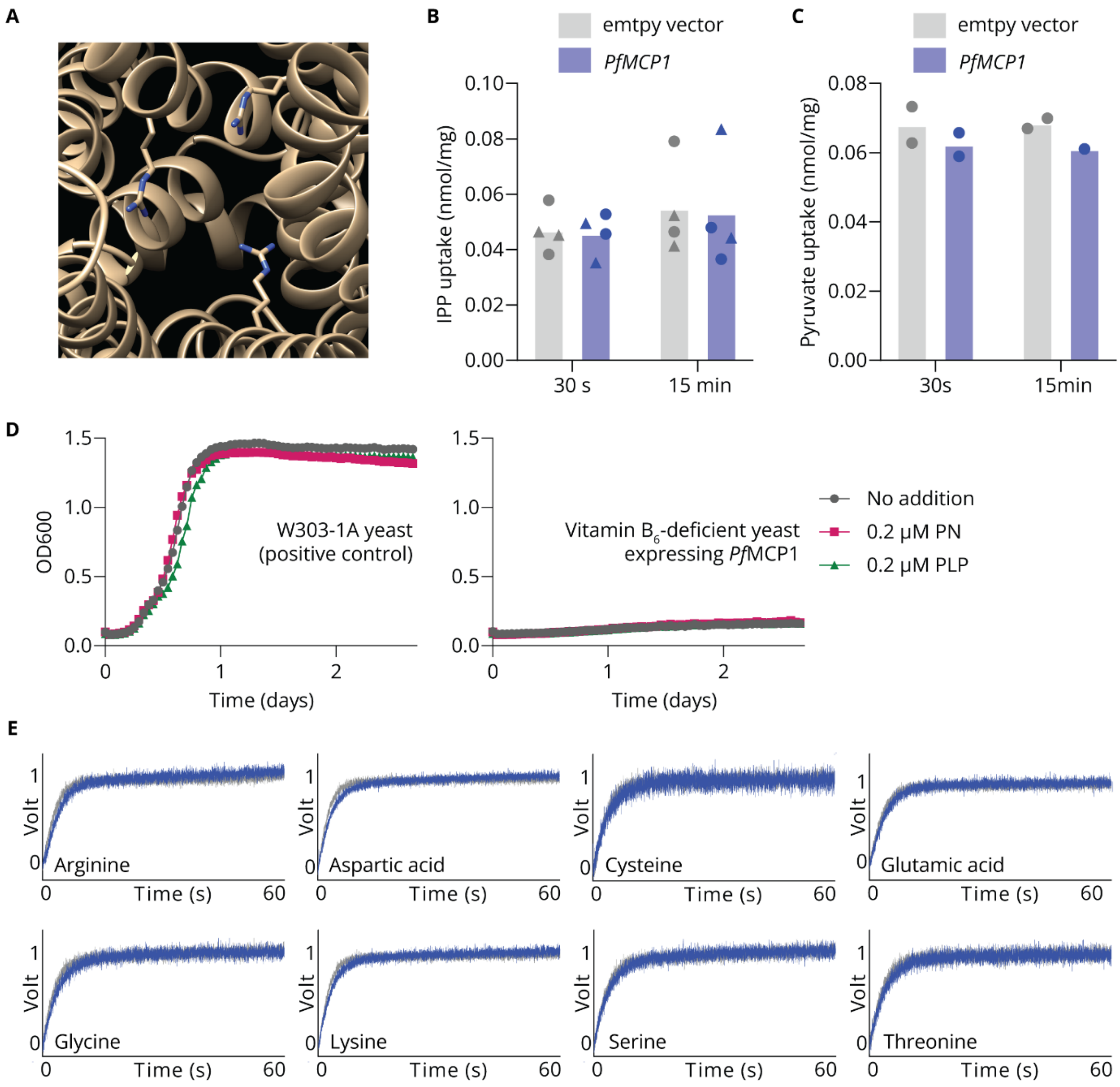


***Figure S5 – Candidate substrates are not taken up by* PfMCP1*-expressing yeast.*** *A) Binding pocket of* PfMCP1*, predicted by AlphaFold 3, with the positively charged side chains of p.Arg153, p.Arg333 and p.Arg416 pointing inwards. B) Uptake of IPP by* PfMCP1*-expressing yeast and negative control as measured by direct transport assays with radiolabeled substrate. Assays were*

*performed in 50 mM TRIS/50 mM HEPES buffer (pH 6.8). Performing the assay at pH 4.8 or pH 8.3, or adding $Na^+$, $K^+$, or $Mg^{2+}$ as a cofactor yielded similar results. Each symbol represents an individual measurement (different symbols for independent experiments). C) Uptake of pyruvate by* PfMCP1*-expressing yeast and negative control as measured by direct transport assay with radiolabeled substrate. Assay was performed in 50 mM TRIS/50 mM HEPES buffer (pH 6.8). Each symbol represents an individual measurement. D) Growth of vitamin $B_6$-deficient yeast expression* PfMCP1 *and positive control in pyridoxin (PN)-free medium with supplemented PN or pyridoxal-5'-phosphate (PLP). E) Stopped-flow light scattering measurements of* PfMCP1*-expressing yeast (blue) or negative control (gray) exposed to a 300 mM inward gradient of various amino acids.*

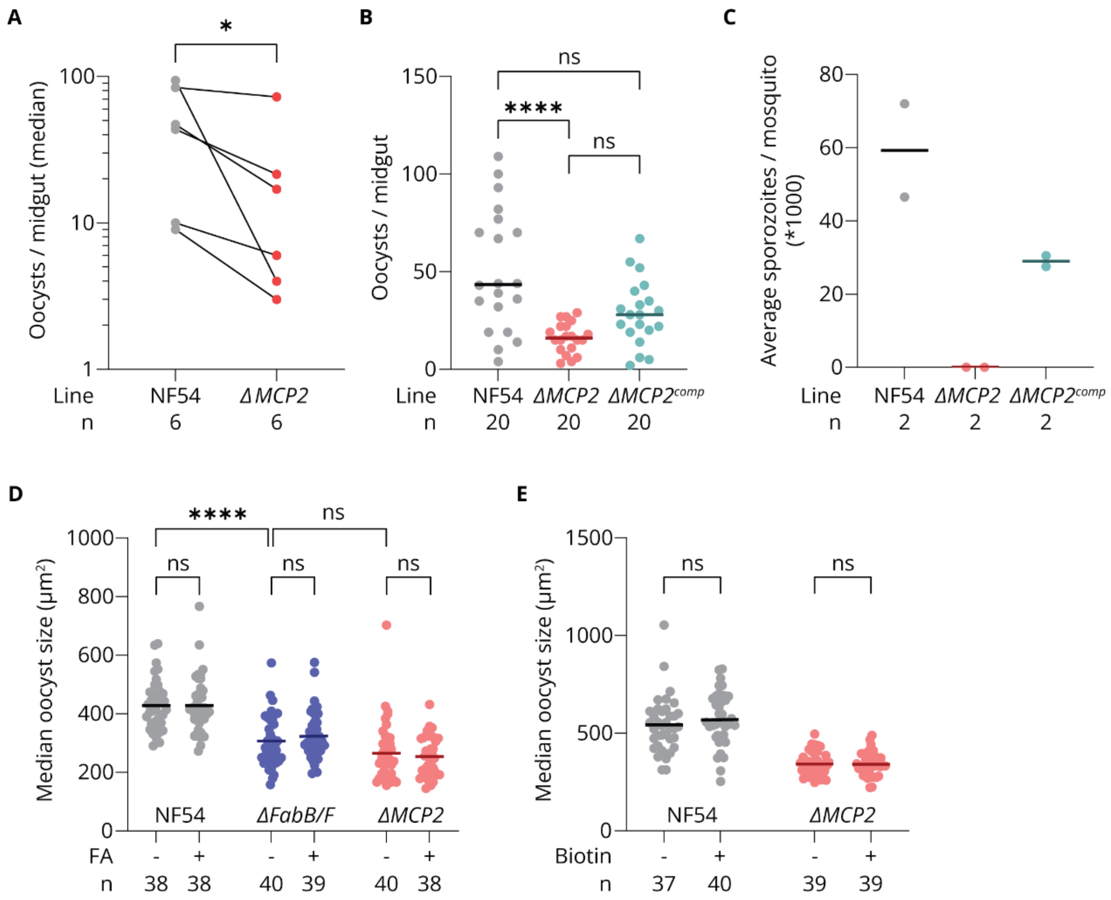


***Figure S6 – Oocyst intensity and size upon* MCP2 *deletion,* MCP2 *complementation, and nutrient supplementation.*** *A) Median oocyst intensity in* Anopheles stephensi *midguts infected with NF54 or* ΔMCP2 *parasites per standard membrane feeding assay. Data points obtained in a single assay are linked. Statistical significance calculated with a Wilcoxon matched-pairs signed-rank test. B) Oocyst intensity in* Anopheles stephensi *midguts after a standard membrane feeding assay with NF54,* ΔMCP2*, or* ΔMCP2$^{comp}$ *gametocytes. significance was calculated with a Kruskal-*

*Wallis test followed by Dunn's test. C) Sporozoite numbers at day 14 after the infectious bloodmeal with NF54 or ΔMCP2 parasites. Salivary glands from 10 mosquitoes were pooled per data point to determine the average number of salivary gland sporozoites per dissected mosquito. D) Median oocyst size per mosquito at day 7 after the infectious bloodmeal with NF54, ΔFabB/F, or ΔMCP2 parasites. After the infectious bloodmeal, mosquitoes were supplemented with 250 μM BSA-coupled fatty acids in 5% glucose water (FA +) or 250 μM BSA in 5% glucose water (FA - ). Mosquitoes with two or less oocysts were excluded from analysis. Differences were assessed with a two-way ANOVA followed by Šídák's test. E) Oocyst size per mosquito at day 7 after the infectious bloodmeal with NF54 or ΔMCP2 parasites. In the biotin+ condition, gametocytes were incubated with 40 μM biotin for 24 h before the bloodmeal, 40 μM biotin was added to the bloodmeal, and the mosquitoes were supplemented with 40 μM biotin in 5% glucose water after the infectious bloodmeal. Each data point represents the median oocyst size in a single mosquito, mosquitoes with two or less oocysts were excluded from analysis. Differences between biotin- and biotin+ conditions for each line were assessed with two-way ANOVA followed by Šídák's test. In all graphs, the lines represent the medians (B, C) or the means (D, E). ns: not significant, *: 0.01 < p < 0.05, ****: p < 0.0001.*

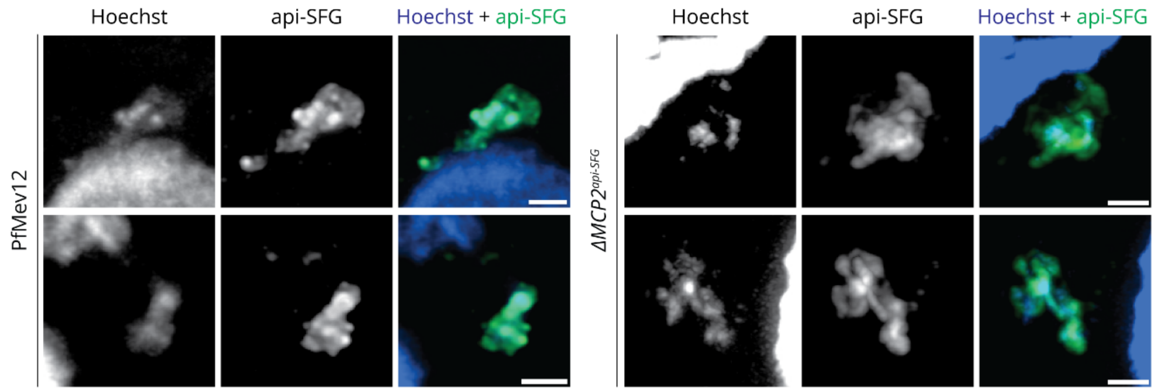


***Figure S7 – Apicoplast DNA in oocysts with and without* MCP2.** *Fluorescence microscopy of oocysts within dissected mosquito midguts, 3 days after and infectious bloodmeal with PfMev12 or ΔMCP2$^{api\text{-}SFG}$ parasites. Images are maximum intensity projections of Z-stacks. Scale bars: 1 μm.*

### Table S1 – Transport assays

| Readout | Advantages | Disadvantages | Used for |
|---|---|---|---|
| Radioactivity | - Very sensitive | - Radiolabeled substrate is expensive<br>- Requires a specialized laboratory | IPP, pyruvate |
| Stopped-flow light scattering | - Cheap<br>- Easy to test many substrates | - Low sensitivity<br>- Requires very high substrate solubility (600 mM) | Amino acids |
| Growth rescue | - Cheap | - Requires deficient yeast strains<br>- Indirect | PLP, pyridoxin |

### Table S2 – Oligonucleotides

| **Oligonucleotides used as primers to generate amplicons for cloning**<br>Sequences that anneal to the template are in capital letters. | | | | | | |
|---|---|---|---|---|---|---|
| ID | Line | Gene | Region | F/R | Sequence | Template |
| ST036 | N/A | N/A | LacZ v1 | F | tttttctcgaggcagagtgagacgactgGCAGCTGGCACGACAGG | pRF0506 |
| ST037 | | | | R | tatatacggccgcgcggtagagacgtgacCGGTCACAGCTTGTCTGTAAG | |
| LA062 | N/A | N/A | LacZ v2 | F | gactctcgagcctagggacatgagacgGCAGCTGGCACGACAGGTTTC | pRF0506 |
| LA063 | | | | R | ctagcggccgctaggacgtcggtaccgtgcagagacgGTCACAGCTTGTCTGTAAGCGGATG | |
| LA058 | FLAG | N/A | HA (Cterm) | F | caggtGGATCCGCTTACCCG | pRF0490 |
| LA059 | | | | R | ctagGAATTCCGAGATCATGTGATTTCTCTTTGTTC | |
| LA137 | FLAG | N/A | FLAG | F | gactggatcccgtctctggatccGGCGGATCAGGTGGATCCG | pRF0740 |
| LA139 | | | | R | ctaggaattccgtctctgtgcCGAGATCATGTGATTTCTCTTTGTTC | |
| LA005 | FLAG | *MCP1* | 5' HR | F | gactggtaccCTAATTCTTTAAGAACATTATCGAATACATCG | NF54 gDNA |

| | | | | | | |
|---|---|---|---|---|---|---|
| LA006 | | | | R | ctagggatccacctgatccgccaaatgctagtgctaacatgatgctcacgaacGATGATAAGGAAACAATTG | |
| LA007 | FLAG | *MCP1* | 3' HR | F | gactgaattcCAATGTATACACAAAATATTTGTAAATCGATAC | NF54 gDNA |
| LA008 | | | | R | ctaggccggcGTGGAGGGTATTTTTTTGTGGTTCATTG | |
| LA127 | FLAG | *MCP2* | 5' HR | F | gactggatcccgtctctagagGATAAATTCTTCTTTTTATTATGGATATCAGTTG | NF54 gDNA |
| LA128 | | | | R | ctaggaattccgtctctatccTTTTGTCAAGATACACATAAGTAAGGTTG | |
| LA129 | FLAG | *MCP2* | 3' HR | F | gactggatcccgtctctgcacGGCATGGCAAAATATAATAAACATG | NF54 gDNA |
| LA130 | | | | R | ctaggaattccgtctctcggtCATTTTAAACGTTTGTTTTGGATGC | |
| LA013 | FLAG | *UMF* | 5' HR | F | gactggtaccGGAAGATGGATGGGTGTACAAAG | NF54 gDNA |
| LA014 | | | | R | ctagggatccacctgatccgccGACAATTTCCTTAACAGGAACTAACC | |
| LA015 | FLAG | *UMF* | 3' HR | F | gactgaattcAGGGAAGAAACAATTTACAAAAAGATTC | NF54 gDNA |
| LA016 | | | | R | ctaggccggcTTACATAAACTGGTAGTACATAATATGGC | |
| LA021 | HA | N/A | HA (Nterm) | F | ggctaccgtctcatgtaCCCGTACGACGTCCC | pRF0205 |
| LA022 | | | | R | ggctaccgtctcaacctcctgatcctccGGCATAATCTGGAACATCGTAAGG | |
| LA023 | HA | *MCP1* | 5' HR | F | ggctaccgtctcaggagCTGGAGAGTATAATAAGAGAGTTGAAAG | NF54 gDNA |
| LA024 | | | | R | ggctaccgtctcatacatTTCTAAATGTTTATATATATATATATATAAATATGTTATACTATG | |
| LA025 | HA | *MCP1* | 3' HR | F | ggctaccgtctcaaggtATGAAAAAAGAGAATACTTCGCTATTGTCATCAAG | NF54 gDNA |
| LA026 | | | | R | ggctaccgtctcaatggCTTACCACTCCTTTGTAATCATAGTGC | |
| LA162 | *Δ* | *MCP1* | 5' HR | F | gactggatcccgtctctagagCTGGAGAGTATAATAAGAGAGTTGAAAG | NF54 gDNA |
| LA163 | | | | R | ctaggaattccgtctctcttaTTCTAAATGTTTATATATATATATATATATAAATATGTTATACTATG | |

| | | | | | | |
|---|---|---|---|---|---|---|
| LA164 | Δ | *MCP1* | 3' HR | F | gactggatcccgtctctgcacCAATGTATACACAAAATATTTGTAAATCGATAC | NF54 gDNA |
| LA165 | | | | R | ctaggaattccgtctctcggtGTGGAGGGTATTTTTTTGTGGTTCATTG | |
| LA125 | Δ | *MCP2* | 5' HR | F | gactggatcccgtctctagagCGGTTGAGATTTCTCCTAAAGTTG | NF54 gDNA |
| LA126 | | | | R | ctaggaattccgtctctcttaCAAAAGTTTTATAAAAATATGTACTTATATACACATG | |
| LA129 | Δ | *MCP2* | 3' HR | F | See LA129 (above) | NF54 gDNA |
| LA130 | | | | R | See LA130 (above) | |
| LA166 | Δ | *UMF* | 5' HR | F | gactggatcccgtctctagagCTATACACCTTTGATTTTTATAGATTAGATAAATG | NF54 gDNA |
| LA167 | | | | R | ctaggaattccgtctctcttaTTTTAAATCATCATTTATTCTGGCTTCC | |
| LA168 | Δ | *UMF* | 3' HR | F | gactggatcccgtctctgcacAGGGAAGAAACAATTTACAAAAAGATTC | NF54 gDNA |
| LA169 | | | | R | ctaggaattccgtctctcggtTTACATAAACTGGTAGTACATAATATGGC | |
| LA092 | $\Delta^{dis}$ | *MCP1* | 5' HR | F | gactaagcttcgtctcagacaGCTTATATTTTGTAAATTAGTATAATATACGAAAGG | NF54 gDNA |
| LA093 | | | | R | ctagggatcccgtctcattacCTGATTTAATATTCCTCCAAAAAAACCAAAAAAAC | |
| LA094 | $\Delta^{dis}$ | *MCP1* | 3' HR | F | gactaagcttcgtctcattagGATGTGGAATAGCATATCCTATCCC | NF54 gDNA |
| LA095 | | | | R | ctagggatcccgtctcagtgcCTAAGTTACTAAAATATTTCTCATCGGAG | |
| LA098 | $\Delta^{dis}$ | *UMF* | 5' HR | F | gactaagcttcgtctcagacaATGGACACAACCGCTATATAATTTAGTAC | NF54 gDNA |
| LA099 | | | | R | ctagggatcccgtctcattacCTTTTTAAATTTCATAAATATATAATGATCATGTAAAAATG | |
| LA100 | $\Delta^{dis}$ | *UMF* | 3' HR | F | gactaagcttcgtctcattagAAAAAAAATGTTACAATTTATATCAGTATCTTATTTATTAG | NF54 gDNA |
| LA101 | | | | R | ctagggatcccgtctcagtgcTCTGAATTTTTTATGGTATCTACTGTTTTATTTTC | |
| LA125 | $\Delta^{comp}$ | *MCP2* | Whole gene | F | See LA125 (above) | NF54 gDNA |
| LA130 | | | | R | See LA130 (above) | |

| | | | | | | |
|---|---|---|---|---|---|---|
| LA188 | $\Delta^{api\text{-}SFG}$ | N/A | Api-SFG | F | gactggtctcttccgCCAGCATGCTATTTGATGAA TTAACTACAC | PfMev gDNA |
| LA189 | | | | R | ctagggtctctatggcgtctctgtgcGGTCATGGATC CTGATATATTTCTATTAGG | |
| LA125 | $\Delta^{api\text{-}SFG}$ | *MCP2* | 5' HR | F | See LA125 (above) | pRF0731 |
| LA126 | | | | R | See LA126 (above) | |
| LA129 | $\Delta^{api\text{-}SFG}$ | *MCP2* | 3' HR | F | See LA129 (above) | pRF0731 |
| LA130 | | | | R | See LA130 (above) | |

**Oligonucleotides that were annealed to use directly for cloning**

For the oligos that were used to clone the 'guide plasmid', the guide sequence is in capitals.

| ID | Line | Gene | Region | F/R | Sequence |
|---|---|---|---|---|---|
| LA147 | FLAG | N/A | N/A | F | gatccgattacaaagatcatgatggagattataaagatcacgatatag attacaaggatcatgactaaa |
| LA148 | | | | R | ctagtttagtcatgatccttgtaatctatatcgtgatctttataatctccat catgatctttgtaatcg |
| LA033 | FLAG | *MCP1* | Guide | F | tattgTGCTAACATGATGCTGACAA |
| LA034 | | | | R | aaacTTGTCAGCATCATGTTAGCAc |
| LA135 | FLAG | *MCP2* | Guide | F | tattgTATCTTGACAAAATAAGGCA |
| LA136 | | | | R | aaacTGCCTTATTTTGTCAAGATAc |
| LA037 | FLAG | *UMF* | Guide | F | tattgTAAGGAAATTGTCTGAGCAA |
| LA038 | | | | R | aaacTTGCTCAGACAATTTCCTTAc |
| LA031 | HA | *MCP1* | Guide | F | tattgTCTATATCTTGATGATAACA |
| LA032 | | | | R | aaacTGTTATCATCAAGATATAGAc |
| LA031 | $\Delta$ | *MCP1* | Guide 1 | F | See LA031 (above) |
| LA032 | | | | R | See LA032 (above) |
| LA033 | $\Delta$ | *MCP1* | Guide 2 | F | See LA033 (above) |
| LA034 | | | | R | See LA034 (above) |
| LA131 | $\Delta$ | *MCP2* | Guide 1 | F | tattgATATACGGCACAGCTGTTGA |
| LA132 | | | | R | aaacTCAACAGCTGTGCCGTATATc |
| LA135 | $\Delta$ | *MCP2* | Guide 2 | F | See LA135 (above) |
| LA136 | | | | R | See LA136 (above) |
| LA035 | $\Delta$ | *UMF* | Guide 1 | F | tattgATAGAATTAATTATTCAATG |
| LA036 | | | | R | aaacCATTGAATAATTAATTCTATc |
| LA037 | $\Delta$ | *UMF* | Guide 2 | F | See LA037 (above) |
| LA038 | | | | R | See LA038 (above) |
| LA102 | $\Delta^{dis}$ | *MCP1* | Guide | F | tattgCAGATTAGTGTCTTATTAGG |
| LA103 | | | | R | aaacCCTAATAAGACACTAATCTGc |

| | | | | | |
|---|---|---|---|---|---|
| LA104 | *$\Delta^{dis}$* | *UMF* | Guide | F | tattgTTAGTTATATCACCAATAGT |
| LA105 | | | | R | aaacACTATTGGTGATATAACTAAc |

**Oligonucleotides used as primers to detect nuclear/apicoplast/mitochondrial genome presence[11]**

| ID | Genome | Gene | F/R | Sequence |
|---|---|---|---|---|
| LA154 | Nuclear | *ldh* | F | GGAGATGTAGTTTTGTTCGATATTG |
| LA155 | | | R | CTTGTAAAGGGATACCACCTACAG |
| LA156 | Apicoplast | *sufB* | F | CATGTAGCTATAGTAGAAATAATAGTAAAAGATTATGG |
| LA157 | | | R | GACTCTGAAATACTTAAACCACGTTGC |
| LA158 | Mitochondrion | *cox1* | F | CTTCATCTTTAAGAATAATTGCACAAGAAAATGTAAATC |
| LA171 | | | R | GTACATATGATGTACCCATACTAAGCTTCC |

**Oligonucleotides used as primers to check genetic modification**

5’ reaction detects 5’ integration; 3’ reaction detects 3’ integration, C (control) reaction detects presence of mother line

| ID | Line | Reaction | F/R | Sequence |
|---|---|---|---|---|
| LA049 | MCP1-FLAG | 5’ | F | CTAAGGATTTCATGGCATATAATGATAGG |
| EB249 | | | R | CGTGATCTTTATAATCTCCATCATGATC |
| EB251 | MCP1-FLAG | 3’ | F | GATCATGATGGAGATTATAAAGATCACG |
| LA050 | | | R | CCTTTTATGGTTTAAATTAAGTCCACACG |
| LA090 | MCP1-FLAG | C | F | GCACTAGCATTTTAATAATATAAAAATGAAC |
| LA050 | | | R | See LA050 (above) |
| LA149 | MCP2-FLAG | 5’ | F | CTCTCTTTTTGGTATGTTAACATTAGATAG |
| EB249 | | | R | See EB249 (above) |
| EB251 | MCP2-FLAG | 3’ | F | See EB251 (above) |
| LA150 | | | R | GAAATAAAACCATGAAGAAATTAAGATACAGTC |
| LA209 | MCP2-FLAG | C | F | CTTGACAAAATAAGGCATGGC |
| LA150 | | | R | See LA150 (above) |
| LA051 | UMF-FLAG | 5’ | F | CAAAATTGCACGAATTCAATAGATAAAAG |
| EB249 | | | R | See EB249 (above) |
| EB251 | UMF-FLAG | 3’ | F | See EB251 (above) |
| LA052 | | | R | GGGTACACACAAATGTATAATAATATGGAG |
| LA091 | UMF-FLAG | C | F | GGAAATTGTCTGAGCAAGGGAAG |
| LA052 | | | R | See LA052 (above) |
| LA040 | HA-MCP1 | 5’ | F | GGGTTTGGTAGAGATGAACCTTTC |
| JV103 | | | R | CTGGAACATCGTAAGGATACGC |
| LA041 | HA-MCP1 | 3’ | F | GCGTATCCTTACGATGTTCCAG |
| LA042 | | | R | CAAAATGGTGAAACGAAATACAACAC |

| | | | | |
|---|---|---|---|---|
| LA040 | HA-MCP1 | C | F | See LA040 (above) |
| LA053 | | | R | GGGAAGTATTCTCTTTTTTCATTTCTAAATG |
| LA040 | *ΔMCP1* | 5’ | F | See LA040 (above) |
| EB028 | | | R | GCACATCCTTATATACATTATGCACATTTATGTATG |
| LA172 | *ΔMCP1* | 3’ | F | CGATTTCTTAATATTGATAAGTATCATGTG |
| LA050 | | | R | See LA050 (above) |
| LA040 | *ΔMCP1* | C | F | See LA040 (above) |
| LA160 | | | R | CACATTAACCATCCACCTAATAAGACAC |
| LA152 | *ΔMCP2* | 5’ | F | GGGCATACAGAAGAGATAATATGTG |
| EB028 | | | R | See EB028 (above) |
| LA172 | *ΔMCP2* | 3’ | F | See LA172 (above) |
| LA150 | | | R | See LA150 (above) |
| LA152 | *ΔMCP2* | C | F | See LA152 (above) |
| LA170 | | | R | CAATATACGGCACAGCTGTTG |
| LA045 | *ΔUMF* | 5’ | F | CACATTGTTTTATAGCATTTTTTTATATATTTCCC |
| EB028 | | | R | See EB028 (above) |
| LA172 | *ΔUMF* | 3’ | F | See LA172 (above) |
| LA052 | | | R | See LA052 (above) |
| LA045 | *ΔUMF* | C | F | See LA045 (above) |
| LA161 | | | R | CTATTGGTGATATAACTAAACAGACACTTG |
| LA040 | *$ΔMCP1^{dis}$* | 5’ | F | See LA040 (above) |
| EB028 | | | R | See EB028 (above) |
| EB155 | *$ΔMCP1^{dis}$* | 3’ | F | CATTTTTCTTTACATAAAAAAACTGTCCAATAAAAAATG |
| LA151 | | | R | CCTATCATTATATGCCATGAAATCCTTAG |
| LA040 | *$ΔMCP1^{dis}$* | C | F | See LA040 (above) |
| LA160 | | | R | See LA160 (above) |
| LA045 | *$ΔUMF^{dis}$* | 5’ | F | See LA045 (above) |
| EB028 | | | R | See EB028 (above) |
| EB155 | *$ΔUMF^{dis}$* | 3’ | F | See EB155 (above) |
| LA153 | | | R | GGCTAGGGTTATCATGTGAGAGAC |
| LA045 | *$ΔUMF^{dis}$* | C | F | See LA045 (above) |
| LA161 | | | R | See LA161 (above) |
| LA152 | *$ΔMCP2^{comp}$* | 5’ | F | See LA152 (above) |
| LA170 | | | R | See LA170 (above) |
| LA127 | *$ΔMCP2^{comp}$* | 3’ | F | See LA127 (above) |
| LA150 | | | R | See LA150 (above) |
| LA152 | *$ΔMCP2^{comp}$* | C | F | See LA152 (above) |

| | | | | |
|---|---|---|---|---|
| EB200 | | | R | GGTGGAACAGTACGAACGCTCC |
| LA152 | *ΔMCP2*$^{api\text{-}SFG}$ | 5’ | F | See LA152 (above) |
| LA143 | | | R | CCATACGATGTGCCTGATTATGC |
| EB250 | *ΔMCP2*$^{api\text{-}SFG}$ | 3’ | F | GAGATATATATGGAGGTGAAAATACCG |
| LA150 | | | R | See LA150 (above) |
| LA152 | *ΔMCP2*$^{api\text{-}SFG}$ | C | F | See LA152 (above) |
| LA170 | | | R | See LA170 (above) |

### Table S3 – Cloning reactions

| **Traditional restriction enzyme digestion and ligation reactions** | | | | | | |
|---|---|---|---|---|---|---|
| ID | Status | Line | Gene | Backbone | Insert | Enzymes |
| pRF0624 | Intermediate | N/A | N/A | pGGAselect | LacZ v1 amplicon | XhoI & EagI |
| pRF0613 | Intermediate | N/A | N/A | pGGAselect | LacZ v2 amplicon | XhoI & EagI |
| pRF0489 | Intermediate | FLAG | *MCP1* | pRF0205 | 5’ HR amplicon | KpnI & BamHI |
| pRF0490 | Intermediate | FLAG | *MCP1* | pRF0489 | 3’ HR amplicon | EcoRI & NgoMIV |
| pRF0592 | Intermediate | FLAG | *MCP1* | pRF0490 | HA amplicon | BamHI & EcoRI |
| pRF0740 | Final | FLAG | *MCP1* | pRF0592 | FLAG oligo | BamHI & SpeI |
| pRF0492 | Intermediate | FLAG | *UMF* | pRF0205 | 5’ HR amplicon | KpnI & BamHI |
| pRF0493 | Intermediate | FLAG | *UMF* | pRF0492 | 3’ HR amplicon | EcoRI & NgoMIV |
| pRF0593 | Intermediate | FLAG | *UMF* | pRF0493 | HA amplicon | BamHI & EcoRI |
| pRF0741 | Final | FLAG | *UMF* | pRF0593 | FLAG oligo | BamHI & SpeI |
| pRF0666 | Intermediate | $\Delta^{dis}$ | *MCP1* | pRF0506 | 5’ HR amplicon | BamHI & HindIII |
| pRF0667 | Intermediate | $\Delta^{dis}$ | *MCP1* | pRF0506 | 3’ HR amplicon | BamHI & HindIII |
| pRF0668 | Intermediate | $\Delta^{dis}$ | *UMF* | pRF0506 | 5’ HR amplicon | BamHI & HindIII |
| pRF0669 | Intermediate | $\Delta^{dis}$ | *UMF* | pRF0506 | 3’ HR amplicon | BamHI & HindIII |

| **Golden Gate Assembly reactions** | | | | | | | |
|---|---|---|---|---|---|---|---|
| ID | Status | Line | Gene | Backbone | Insert 1 | Insert 2 | Insert 3 |
| pRF0733 | Final | FLAG | *MCP2* | pRF0624 | 5’ HR amplicon | FLAG amplicon | 3’ HR amplicon |
| pRF0495 | Final | HA | *MCP1* | pGGAselect | 5’ HR amplicon | HA (Nterm) amplicon | 3’ HR amplicon |
| pRF0742 | Final | *Δ* | *MCP1* | pRF0624 | 5’ HR amplicon | pRF0508 | 3’ HR amplicon |

| | | | | | | | |
|---|---|---|---|---|---|---|---|
| pRF0731 | Final | $\Delta$ | *MCP2* | pRF0624 | 5’ HR amplicon | pRF0508 | 3’ HR amplicon |
| pRF0743 | Final | $\Delta$ | *UMF* | pRF0624 | 5’ HR amplicon | pRF0508 | 3’ HR amplicon |
| pRF0675 | Final | $\Delta^{dis}$ | *MCP1* | pRF0613 | pRF0666 | pRF0674 | pRF0667 |
| pRF0676 | Final | $\Delta^{dis}$ | *UMF* | pRF0613 | pRF0668 | pRF0674 | pRF0669 |
| pRF0783 | Final | $\Delta^{comp}$ | *MCP2* | pRF0624 | Whole gene amplicon | N/A | N/A |
| pRF0784 | Final | $\Delta^{api\text{-}SFG}$ | *MCP2* | pRF0624 | 5’ HR amplicon | Api-SFG amplicon | 3’ HR amplicon |
